# Intracellular accumulation of AAV6 genomes governs microglia reactivity and the inflammatory response in retinal gene therapy

**DOI:** 10.64898/2026.10.09.757886

**Authors:** Jean-Baptiste Ducloyer, Allwyn Pereira, Joanna Demilly, Estelle Toublanc, Lyse Libeau, Bérangère Robert, Anthony Mellet, David Augereau, Léa Comtesse, Thibaut Larcher, Alix Lebreton, Paul Socheleau, Marion Grard, Malo Journou, Célia Couzinié, Véronique Blouin, Michel Weber, Guylène Le Meur, Caroline Le Guiner, Oumeya Adjali, Therese Cronin

## Abstract

Adeno-associated viral (AAV) vectors possess immense potential for gene therapy applications. However, microglia can mount an immune response against AAV vectors. Yet, microglia are therapeutic targets since they play a driving role in the progression of neurodegenerative diseases. Thus, it is critical to understand the cellular processes driving the microglia-mediated immune response against AAV vectors.

To this end, we modelled AAV6 induced inflammation upon retinal gene therapy in minipigs. Upon AAV6 administration, the IBA1+ cells demonstrated a morphological transition to the amoeboid form, indicating microglial reactivity. We also observed the invasion of peripheral immune cells in the retina. To elucidate the underlying molecular mechanisms, we transduced human iPSC derived microglia-like cells (iMGLs) *in vitro* with AAV6 and AAV8 respectively. Only AAV6 transduced iMGLs exhibited morphological features and a transcriptomic signature of microglial reactivity. The magnitude of intracellular AAV6 viral genomes was significantly higher as compared to the AAV8 condition. The elevated levels of intracellular AAV6 genomes induced a DNA damage response in AAV6-transduced iMGLs, which is a key feature of neuroinflammation-linked microglial reactivity.

Thus, our work highlights the impact of intracellular viral genome accumulation on microglial reactivity and its implications for retinal gene therapy associated inflammation.

## Introduction

Since 2018, retinal gene therapy has been in the clinic in Europe, with patients treated with Luxturna^®^, an adeno-associated virus serotype 2 (AAV2) vector carrying the human RPE65 transgene ^1,2^. The success of Luxturna^®^ has paved the way for the development of numerous gene therapy candidates to treat an array of inherited retinal diseases (IRDs) and complex retinal diseases such as age-related macular degeneration (AMD), of which many have since undergone or are undergoing clinical trials ^3–5^. While Luxturna^®^ represents gene therapy’s earliest success, studies have reported ocular inflammation, including chorioretinal inflammation and atrophy, in up to 50% of the patients receiving the clinical-grade gene therapy AAV products ^6,7^. The umbrella term “Gene Therapy Associated Uveitis (GTAU)” has come to describe this intraocular immune response that can arise following retinal gene therapy. However, multiple factors such as the surgical procedure, the viral vector serotype and its dose or the retinal state can contribute to the generation of this inflammatory reaction and is an area of active investigation ^8^. Both the acute and chronic form of GTAU will induce irreparable retinal damage and impairment in visual function. Thus, it is critical to identify the cellular mechanisms contributing to this form of inflammation to improve the safety and efficacy of retinal gene therapy.

In the retina, microglia, retinal pigment epithelium (RPE) and Muller cells are capable of recognizing pathogen and damage associated molecular patterns and initiating a pro-inflammatory response ^9^. Microglia are tissue resident macrophages of the central nervous system (CNS), where they play a critical role in the maintenance of homeostasis by performing functions such as tissue surveillance for molecular patterns of injury or infection, phagocytosis, synaptic pruning and the secretion of cytokines. Depending upon the environmental signals, microglia will undergo cell state transitions, characterized by distinct morphological, molecular and functional properties, to enable the CNS to cope with the loss of homeostasis ^10^. Genetic variations in genes associated with a higher risk of developing AMD have been shown to be expressed in microglia, thus making it a pertinent therapeutic target for gene therapy applications ^11^. However, numerous studies have demonstrated a microglia-mediated inflammatory response to AAV vectors ^12–16^. Recently, the employment of single cell and spatial transcriptomics have demonstrated that myeloid cells are recruited to the site of AAV delivery, secrete chemokines which in turn recruit CD8 effector memory cells, ultimately driving a type 1 cell-mediated antiviral response ^14^. This was in line with a similar mechanism of GTAU observed in mice upon subretinal delivery of AAV vectors^12^. Nevertheless, in these studies, microglia were not the intended cellular target for the AAV vectors employed. While, it is established that microglia are resistant to robust transgene expression by AAV vectors, the transient presence of AAV vector genomes in microglia has been reported ^15,17^. Thus, there is a need to elucidate the cell autonomous mechanisms of AAV-induced microglial reactivity and its involvement in GTAU.

Among the AAV serotypes, AAV6 has demonstrated potential for the transduction of microglia and macrophages due to its dual glycan-binding capacity, (for both N-linked sialic acid and Heparan sulfate proteoglycan). Thus, the AAV6 serotype has gained interest in the gene therapy community for CNS applications in recent years and studies have demonstrated that capsid engineering of AAV6 can improve its ability to transduce microglia ^18–21^. However, the risk of inducing microglial reactivity upon AAV6 transduction remains unknown. Therefore, we were interested in modelling the immune response upon AAV6 transduction of microglia, especially concerning the risk of inducing GTAU.

The choice of the animal model for GTAU characterization is critical since rodent models used for preclinical studies show a muted immune response compared to non-human primate (NHP) models ^22^. Moreover, for macular-directed gene transfer, geographic activation of immune cells specifically due to transduction of macular cones cannot be tested in rodents as they lack a cone-rich macular region. NHPs offer the closest anatomy to humans, but their use for pilot exploratory purposes is ethically and financially sensitive. The use of minipigs as a model organism for testing AAV vector safety offer several advantages: presence of a cone-rich visual streak that is functionally similar to the macula, similar size of eyeball structures, absence of *tapetum lucidum* and a well-described surgical approach of the subretinal space that is identical to that in humans ^23–29^.

In this study, we modeled retinal immune cell activation after subretinal injection of AAV6. A subretinal injection of 4.5E11 vg/eye of AAV6 carrying a ubiquitous CAG promoter driving EGFP expression was used to assess the inflammatory response to a high AAV dose in the outer retina. We employed human iPSC derived microglial-like cells (iMGLs) to further characterize the cell autonomous effect of AAV transduction *in vitro* and compare it with AAV8, an AAV serotype that has been demonstrated to induced microglia reactivity *in vivo* ^30^. Our study revealed that the intracellular accumulation of AAV6 genomes in iMGLs triggers a nucleic acid sensing mechanism that mediates microglial reactivity and drives the inflammatory response.

## RESULTS

### Study Design

Vitrectomy and subretinal injection were performed under general anesthesia in the right eye (OD) and the left eye (OS) of Yucatan minipigs with AAV6-CAG-EGFP (n = 3). Minipigs injected with saline (n = 2) in both eyes and an uninjected minipig (n = 1) were used as controls. The OD and OS of one minipig was injected with AAV8-CAG-EGFP (n = 1), since AAV8 has been shown to induce microglia activation *in vivo* ^30^ (Fig. 1A). The surgeries proceeded with no complications. Subsequently, a range of *in vivo* and post-mortem analyses were conducted to characterize the immune response and retinal function (Fig. 1A and Supplementary table 1).

**Figure 1:**
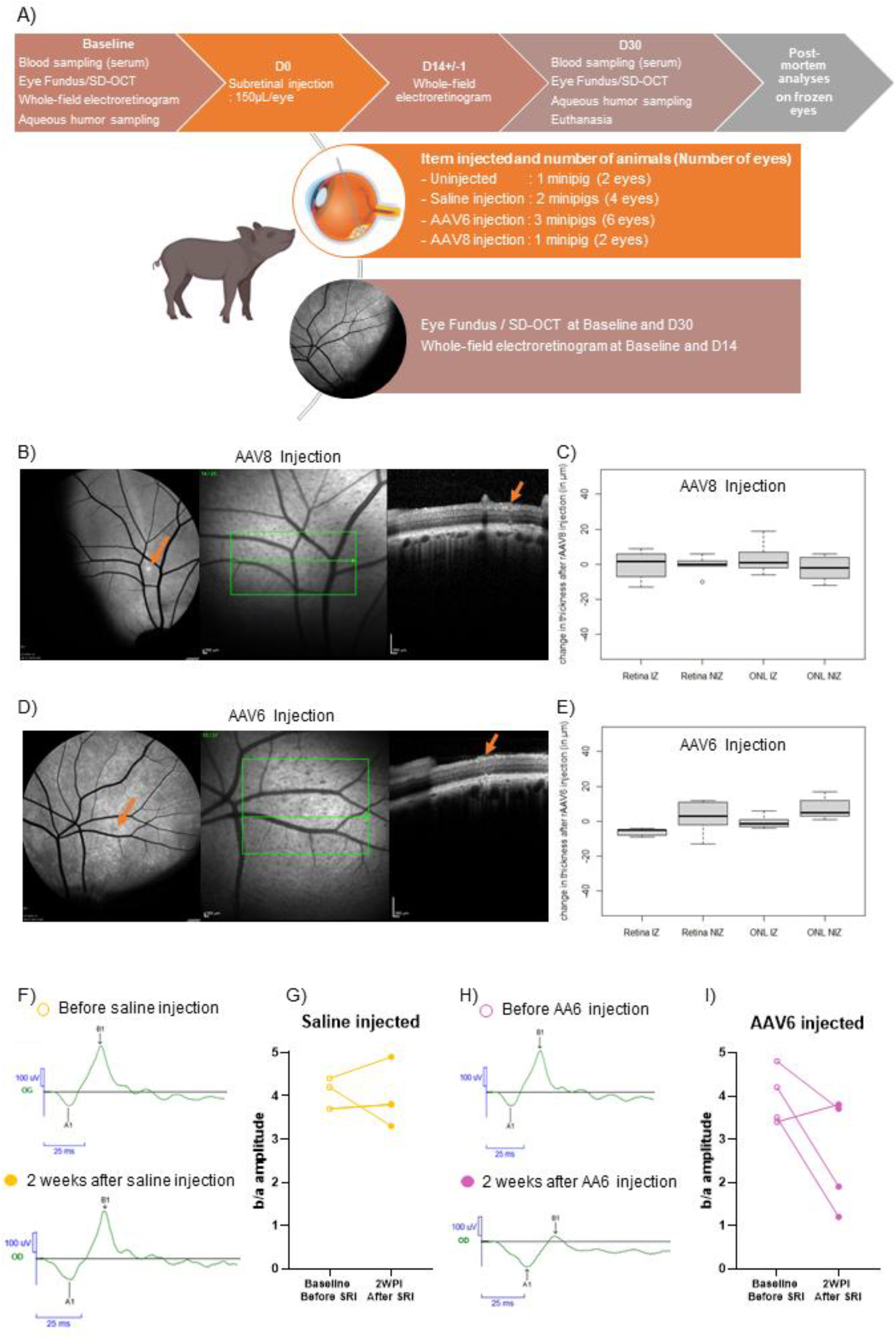
Subretinal injection of AAV6 induces gene-therapy associated inflammation in minipig retina. A) Graphical representation of the *in vivo* study plan. B) Fundus photographs of AAV8 and (D) AAV6 injections with OCT (optical coherence tomography) of retinas at the injection site (shown by an orange arrow). (C and E) The total retinal thickness (RT) and outer nuclear layer thickness (ONL) was measured in the injected zone (IZ) and non-injected zone (NIZ) between baseline and 1 month after surgery in AAV8 and AAV6 injected retinas, respectively. n = 2 eyes for AAV8 and AAV6 injected conditions (F and H) Electroretinograms (ERG) were performed before and two weeks after surgery for the two saline-injected and AAV6-injected retinas, respectively. Representative electroretinogram curves obtained with flashes of 3.0 cd.s/m² at 10 seconds intervals are shown. (G and I) The b/a ratio is plotted for both (G) saline-injected eyes, represented by empty yellow circles before injection and filled yellow circles after injection, and (I) AAV6-injected eyes are represented by empty magenta circles before injection and filled magenta circles after injection. SRI = subretinal injection, D = day, WPI = weeks post-injection. n = 3 eyes for saline-injected condition and n = 4 eyes for AAV6-injected condition.

### Ocular inflammation with impairment of retinal function in AAV6-injected retinas

The postoperative period was normal for all animals without any immediate sign of inflammation during the first week. At the one-month follow-up, Spectral Domain Optical coherence tomography (SD-OCT) showed a moderate increase in autofluorescence in the injected zone (IZ) of both eyes of the AAV8-injected minipig and in one AAV6-injected minipig, as evidenced by confocal scanning laser ophthalmoscopy and shown in EGFP-mediated green fluorescence by color fundus photographs (Supplementary Fig. S1a-d). The retinotomy site, through which the injection was performed, had healed and was no longer noticeable (Fig. 1B and 1D and Supplementary Fig. S1e-j) and the OCT did not show any structural difference between IZ and NIZ (Fig. 1C and 1E).

However, for the remaining two minipigs injected with AAV6-CAG-EGFP, autofluorescence imaging at 1 month was not possible due to clouding of the ocular medium. At day 8 post-surgery, the right eyes already exhibited clinical signs of inflammation: conjunctiva redness, cloudy media, iridocrystalline synechiae and signs of vitritis. Two weeks after surgery, both eyes showed severe intra-ocular inflammation. Whole-field electroretinogram (ERG) was performed on the eyes from saline and AAV6 injected minipigs under baseline conditions (3 weeks before surgery) and day 14 post-surgery to assess global retinal function (Fig. 1A). In the saline-injected retinas, the recordings were reproducible and similar between the pre-operative and post-operative periods (Fig. 1F-G). By contrast, the ERG recordings in the AAV6-injected retinas with overt signs of ocular inflammation showed alteration of function in both eyes for one pig and in just the right eye of the second pig. (Fig. 1H-I). This change involved an increase in the a-wave showing initial photoreceptor response while the post-synaptic b-wave was reduced (Fig. 1H). Changes in inner retinal cell function or circuitry may account for this reduction in b/a ratio between the pre-operative and post-operative periods.

Two of the three AAV6-injected minipigs exhibited ocular inflammation with signs of discomfort due to apparent loss of vision, which involved turning in circles and lethargy. Therefore, general anesthesia for retinal examination and euthanasia was performed earlier than intended, at four weeks post-injection. The cloudy media of the eye indicating vitritis and synechiae prevented end-stage imaging in these two AAV6-injected minipigs, thus reducing the sample size available for the post-mortem analyses and making the *in vivo* results exploratory in nature. Thus, while the surgical procedure employed for subretinal injection of AAV vectors was well tolerated by the animals, 2 out of the 3 minipigs injected with AAV6 exhibited signs of retinal inflammation and impaired retinal function, with the earliest indication of toxicity at 8 days post-injection. Sub-retinal injection of saline or AAV8 neither impaired retinal structure nor did it induce any apparent signs of inflammation at any time-point.

### AAV6 subretinal injection induces migration and morphological changes of IBA1+ cells

We investigated the role of the resident innate immune cells in mediating retinal inflammation upon AAV injection. Mononuclear phagocytes such as macrophages and retinal microglia are the main immune cell types present in the healthy retina. These innate immune cells express pattern recognition receptors such as TLR9 that are capable of recognizing pathogen associated molecular patterns such as unmethylated CpG motifs, which contributes to the initiation of an inflammatory response ^31^ Thus, we performed immunohistochemical analysis to examine the localization of IBA1+ cells and the photoreceptor cell layer of the injected retinas. Retinal cryosections were stained with IBA1 and PNA-lectin to identify IBA1+ cells and external segments of photoreceptors, respectively. The saline injected retinas showed no EGFP-expression, as well as normal outer segment organization (as evidenced by PNA-lectin staining) and absence of IBA1+ cells in the outer retina (Fig. 2A). The AAV8 injected retinas revealed bright EGFP-expression, with well-organized retinal layers, intact external segments of photoreceptors and the IBA1+ cells were mostly localized in the ganglion cell layer (GCL) (Fig. 2A). EGFP expression was found predominantly in the ONL with some staining in the inner retina and low levels of expression extending to the ganglion cell layer. AAV6-injected retinas showed EGFP expression exclusively in the ONL. One of the AAV6-injected minipigs that showed no obvious signs of retinal inflammation and had intact retinal layers and normal PNA lecting staining. However, the retinas of the two AAV6 injected minipigs, that experienced ocular inflammation, had visible disruption of the photoreceptor layer, with complete absence of PNA-lectin staining. This was accompanied by the subretinal infiltration of IBA1+ cells, including in the EGFP positive area (Fig. 2A). We quantified the number of IBA1+ cells in the injected zone (subretinal zone). In the AAV6-injected retinas, we detected 61.3 ± 34.9 IBA1+ cells in the injected zone. In the saline and AAV8-injected retinas, we detected 14 ± 6.4 and 23.5 ± 6.3 IBA1+ cells, respectively (Fig. 2B). The migration of IBA1+ cells to the ONL and subretinal space is a hallmark of retinal inflammation caused during disease and injury ^11,32,33^. Furthermore, subretinal injection of AAV vectors has been shown to induce the microglia activation and migration to the ONL ^16^.

**Figure 2:**
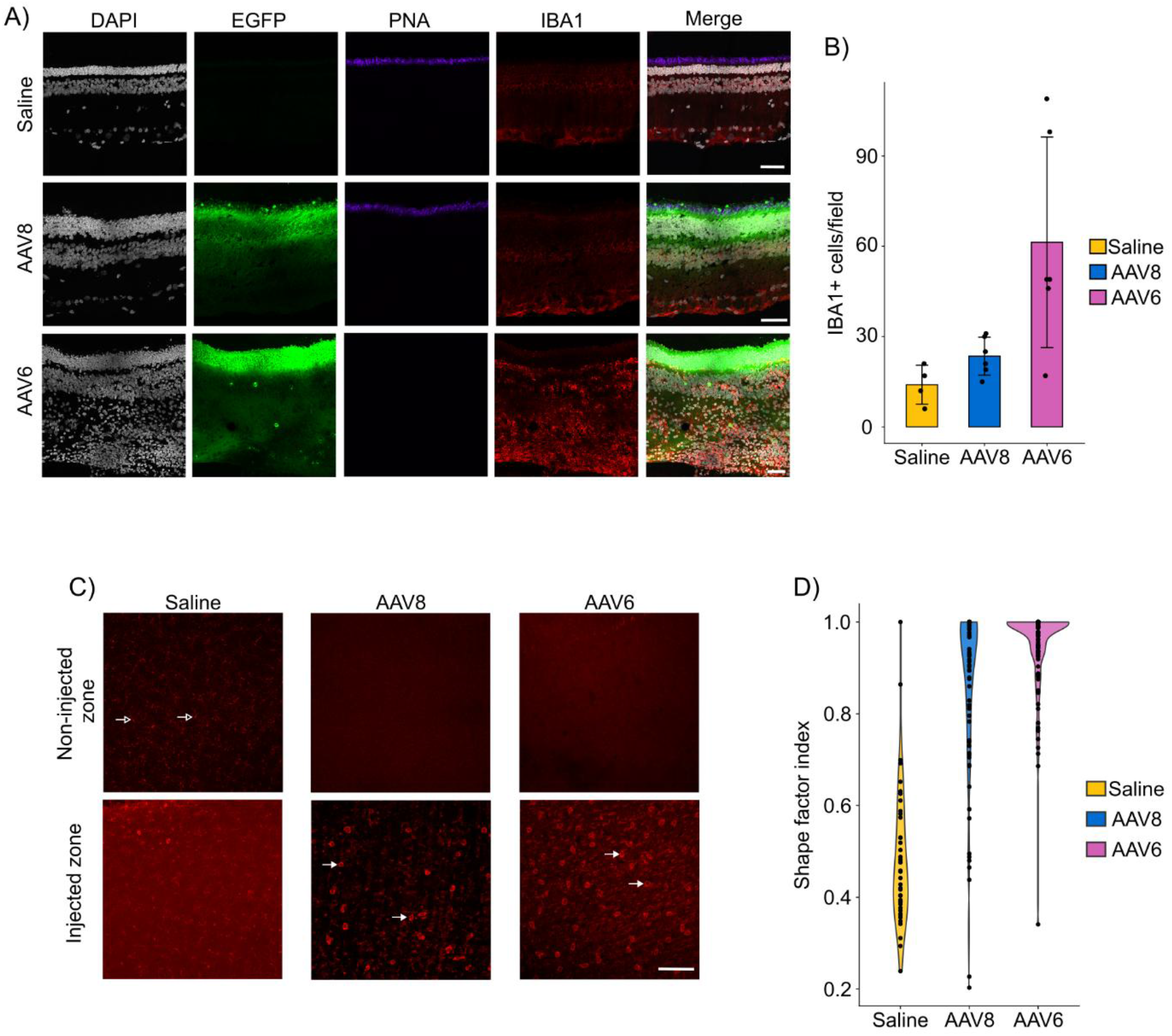
Subretinal injection of AAV6 induces the migration of amoeboid IBA1+ cells into the injected zone. A) Representative images from confocal microscopy (60x) of retinal sections from saline-injected (n=2), AAV8-injected (n=2) and AAV6-injected (n=2) condition. Green corresponds to EGFP expression, red to IBA1 expression, purple to PNA-lectin expression and grey to DAPI expression. Scale bar = 50µm. B) Histogram depicting the quantification of IBA1+ cells in the subretinal zone of saline, AAV8, and AAV6 injected retinas, respectively. n = 1 eye for each experimental condition. Data are plotted as mean ± standard deviation. C) Representative micrographs depicting the effect of saline, AAV8 and AAV6 injections on IBA1+ cell morphology in the peripheral non-injected zone (NIZ) and injected zone (IZ); Filled arrowheads depict IBA1+ cells with an amoeboid morphology and empty arrowheads depict IBA1+ cells with a ramified morphology; scale bar = 100μm. D) Violin plot showing the distribution of the shape factor index of IBA1+ cells in the injected zones of flat mounted retinas with n = 1 eye for each biological condition. A count of 45 cells, 78 cells and 107 cells, for the conditions, saline-injected, AAV8-injected, and AAV6-injected, was recorded, respectively.

In the healthy retina, IBA1+ cells exhibit a ramified morphology, which is a characteristic of homeostasis, while in the context of disease and injury, these cells transition to a reactive state, which is associated with an amoeboid morphology ^10,34^. Thus, we examined the morphology of the IBA1+ cells in the saline and AAV-injected retinas. In the retina injected with saline solution, we observed a network of ramified IBA1+ cells. In contrast, the IBA1+ cells in both AAV6 and AAV8 injected retinas demonstrated a loss of their ramified morphology (Fig. 2C). To assess if this loss in ramification was accompanied by a shift to an amoeboid shape, we employed a semi-automated algorithm to segment and measure the morphological parameters of individual IBA1+ cells. We calculated the shape factor index on the basis of these parameters, where cells with a round morphology possessed a shape factor closer to 1. We confirmed that the shape factor of IBA1+ cells in the AAV6 and AAV8 injected retinas approaches 1, while the saline injected retina exhibited a heterogeneous distribution of shape factors (Fig. 2D).

Thus, subretinal administration of both AAV6 and AAV8 vectors leads to the polarization of IBA1+ cells towards a morphology associated with the reactive cell state. However, the increased inflammatory response in AAV6 injected retinas was correlated by an elevated presence of these activated IBA1+ cells in the outer retinal layer and loss of PNA lecting staining in the photoreceptor cell layer.

### AAV6 subretinal injection is associated with peripheral immune cell invasion

Our results demonstrated that the subretinal administration of AAV vectors led to the polarization of IBA1+ cells to an amoeboid, reactive morphology and their migration into the outer retinal layer, with AAV6-injected retinas displaying hallmark features of retinal inflammation upon AAV delivery ^12–14,16^. Thus, we sought to further define the immune profile of the retina upon subretinal administration of AAV vectors by performing immunohistochemistry for markers of peripheral immune cells. To this end, we stained retinal sections with anti-CD3, CD4, and CD8 antibodies to identify T-cells, anti-CD79 antibody to identify B-cells and anti-CD163, CD14 antibodies to identify monocyte/macrophages cells. We confirmed the absence of B-cells, T-cells and monocyte/macrophages in the saline-injected- and AAV8-injected retinas (Fig. 3A). However, in the AAV6-injected retinas, we detected the presence of CD79+ B cells, CD8+ T-cells and CD163+ macrophage/monocyte cells (Fig. 3A). The aqueous humor of the anterior chamber was assayed for the presence of diverse cytokines pre- and post-injection. IL-8 levels were found to be upregulated in AAV6-injeted retinas (Fig. 3C). The levels of IFNγ did not show a visible pattern upon AAV administration (Fig. 3B). We profiled other cytokines such as GM-CSF, IL-1α, IL-1β, IL-1RA, IL-2, IL-4, IL-6, IL-12, IL-18, TNFα and IL-10. However, the levels of these cytokines were below the detection limit.

**Figure 3:**
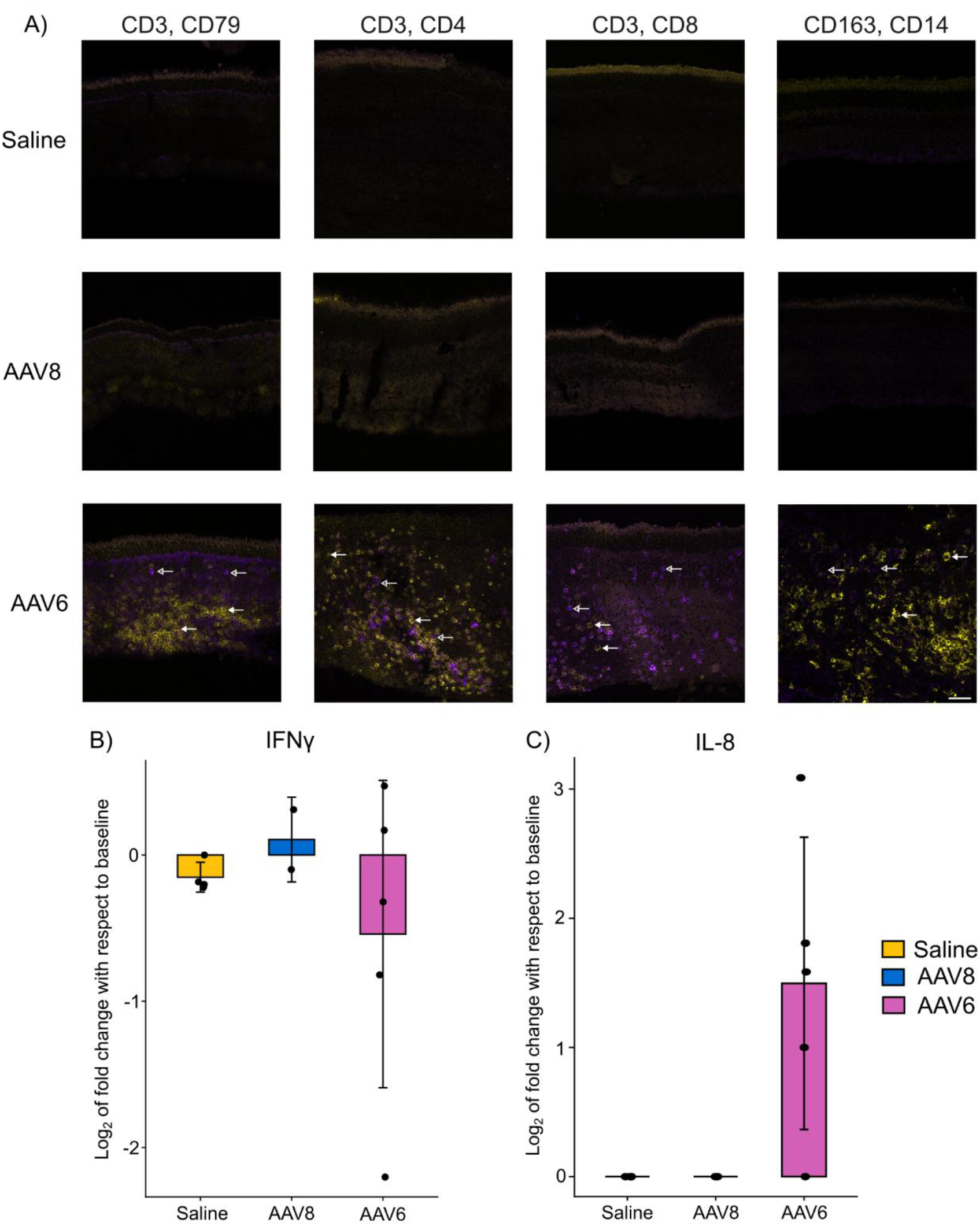
Subretinal injection of AAV6 leads to immune cell infiltration and inflammatory cytokine secretion in the retina. A) Representative images from confocal microscopy (20x) of retinal sections from saline-injected (n=2 eyes), AAV8-injected (n=2 eyes), AAV6-injected (n=4 eyes). In the first column, yellow corresponds to CD3+ cells and purple corresponds to CD79+ cells; in the second column, yellow corresponds to CD3+ cells and purple corresponds to CD4+ cells; in the third column, yellow corresponds to CD3+ cells and purple corresponds to CD8+ cells; in the fourth column, yellow corresponds to CD163+ cells and purple corresponds to CD14+ cells. Filled arrowheads depict the yellow cells and the empty arrowheads depict purple cells in AAV6-injected retinas. Scale bar = 50µm. B-C) Histograms depicting the level of fold change (log_2_) of the cytokines, IFNγ and IL-8, as measured by the Luminex-based multiplex cytokine assay in samples of the anterior chamber fluid. The baseline refers to the cytokine levels before the subretinal injection. The number of samples for each condition are as follows: saline-injected (n = 4 eyes), AAV8-injected (n = 2 eyes) and AAV6-injected (n = 5 eyes). Data are plotted as mean ± standard deviation.

Thus, our data demonstrated that the subretinal administration of AAV6 induces an infiltration of innate and adaptive immune cells in the retina, thus implying their role as active participants in the AAV6-induced inflammatory response.

### Influence of AAV serotype on IBA1+ cell transduction

Next, we examined if the polarization of IBA1+ cells towards the inflammatory state was driven by the internalization of AAV vectors. While no EGFP expression was directly observed in IBA1+ cells, these cells have been shown to be highly permissive to AAV cellular entry ^15^. Thus, we performed RNAscope on histological sections to examine the localization of *IBA1* mRNA, AAV vector DNA (CAG promoter probe) and *EGFP* mRNA. In corroboration with migration of IBA1+ cells to the photoreceptor layer upon inflammation (Fig. 2A-B), we observed strong localization of *IBA1* mRNA (red dots) in the outer retinal layer of AAV6 injected retinas (Fig. 4E and 4G). In the AAV6-injected retina with no ocular inflammation (Minipig #6.1), very few cells exhibited co-localisation of *IBA1* mRNA with CAG promoter DNA probe (violet dots) (Fig. 4F). On the other hand, numerous cells in the AAV6-injected retina with ocular inflammation (Minipig #6.2) exhibited co-localisation between *IBA1* mRNA and CAG promoter DNA probe (Fig. 4H). While *IBA1* mRNA positive cells were observed in the saline-injected and AAV8-injected retinas, none of these cells showed co-localisation with the CAG promoter DNA probe (Fig. 4A-D).These results suggested that AAV6 vectors are capable of entering into IBA1+ cells. However, it remained possible that the CAG promoter DNA probe signal in IBA1+ cells was a result of phagocytosis of transduced photoreceptor cells. Given the extremely low proportion of retinal IBA1+ cells, it was challenging to dissect the cell autonomous and non-cell autonomous effects of AAV transduction and its effect on cell state dynamics in a complex *in vivo* environment.

**Figure 4:**
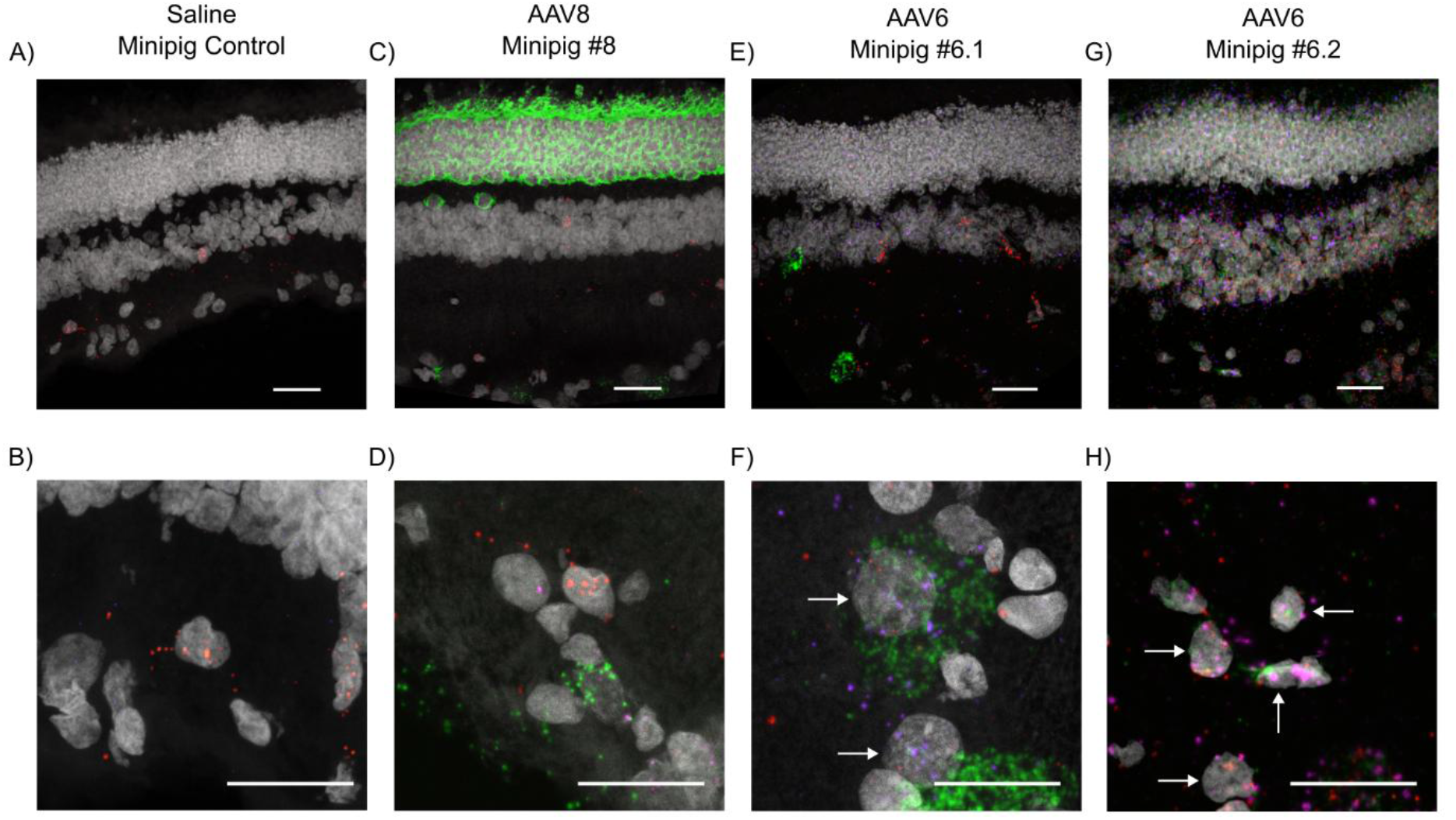
Transduction of IBA1+ cells *in vivo* by AAV6. A-H) Confocal images of histological sections of retinas of saline-injected (minipig control), AAV8-injected (minipig #8) and AAV6-injected animals (minipigs #6.1 and #6.2. Green dots corresponds to RNA from the *EGFP* transgene, red dots to *IBA1* mRNA and violet dots to AAV vector DNA. Scale bar = 50μm. Filled arrowheads depict the co-localization of *IBA1* mRNA dots with AAV vector DNA dots. n = 2 eyes for the conditions, saline and AAV8, n = 4 eyes for the AAV6 condition.

### iMGLs are refractory to transgene expression by AAV vectors

To overcome these limitations, we employed iMGLs, which were differentiated from two independent healthy, human induced pluripotent stem cell (iPSC) lines to model the effect of AAV transduction on microglia reactivity ^35^. We employed flow cytometry analysis to quality control the differentiation process. Our analysis revealed the loss of the pluripotency marker (OCT4) and the gain of a homeostatic microglial marker (P2RY12) in the iMGLs (Supplementary Fig. S2A-F). Prior to employing the iMGL model to elucidate the effect of AAV transduction on microglial reactivity, we aimed to test the pertinence of these cells for modelling aspects of microglial biology. The challenge of achieving robust transgene expression from AAV vectors in microglia has been well documented ^15,17,36^. Thus, we examined if iMGLs are resistant to transgene expression by AAV vectors. To this end, we transduced iMGLs with AAV6 and AAV8 respectively at a MOI of 10^5^ and the cells were immunostained for IBA1 and EGFP expression at 24 h, 48 h 72 h post-transduction. At 24 h post transduction, we observed a very low proportion of EGFP expressing cells, regardless of the AAV serotype. The mean percentage of EGFP expressing cells for AAV6 was 0.4% (Fig. 5A-B). At 72 h post-transduction, no increase in the mean level of EGFP+ cells was observed for AAV8 (0.0% at 24 h versus 0.025% at 72 h). However, a slight increase in the mean of the EGFP+ cells was observed at 72 h post transduction with AAV6, even if the overall transduction efficiency remained low (0.4% at 24 h versus 3.975% at 72 h) (Supplementary Fig. S3A and Fig. 5C-D).

**Figure 5:**
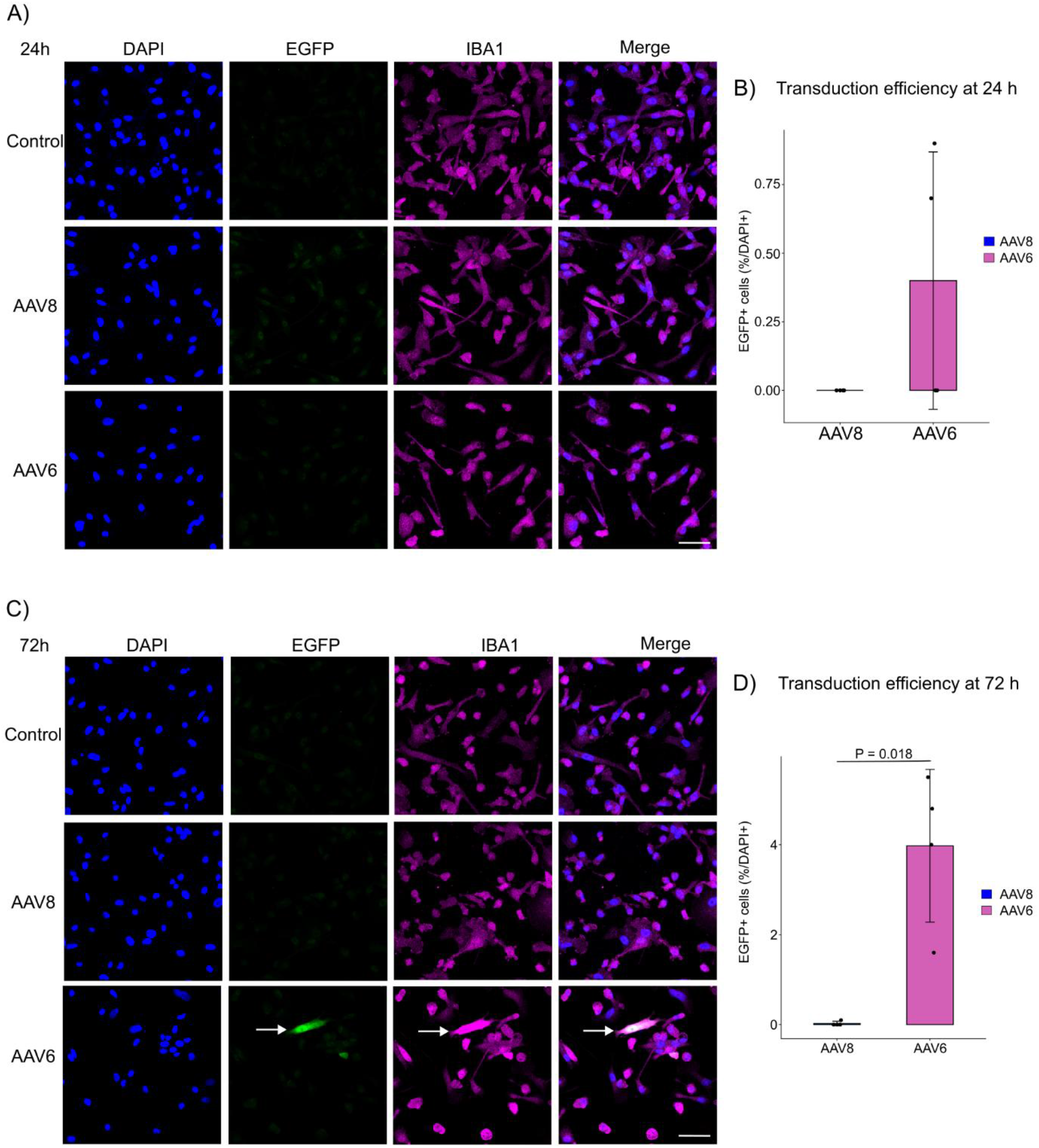
iMGLs are refractory to robust transgene expression by AAV vectors. A and C) Representative micrographs of iMGLs transduced with AAV serotypes, AAV8 and AAV6, respectively at 24 h and 72 h, respectively. Filled arrowheads depicts a EGFP-positive iMGL. Scale bar 50 µm. n = 4 biological replicates. B and D) Histograms depicting the percentage of EGFP+ cells among DAPI+ cells (y-axis) over the AAV serotype (x-axis) at 24 h and 72 h, respectively. Data are plotted as mean ± standard deviation; each dot represents an individual biological replicate (n = 4 biological replicates). Statistical significance was calculated using the Kruskal-Wallis test with Dunn’s multiple comparison test.

These results suggested that neither of the AAV serotype (AAV6 and AAV8) are capable of driving robust transgene expression in iMGLs *in vitro*. Nevertheless, the proportion of EGFP+ cells was significantly higher in AAV6 transduced iMGLs as compared to AAV8 at 72 h post transduction, thus suggesting a serotype dependent effect on AAV cellular entry and transduction (Fig. 5D).

### AAV6 transduction of iMGLs induces morphological and molecular alterations associated with microglial reactivity

Next, we assessed the morphological dynamics of iMGLs upon AAV transduction. To this end, we employed a semi-automated algorithm to segment iMGLs cell bodies and measure their morphological parameters ^37^. We validated the method by assessing the morphology dynamics of iMGLs upon acute ATP treatment. The extracellular concentration of ATP increase upon tissue damage and this acts as a strong chemotactic signal for microglia, leading rapid process extension ^38^. Exposure of microglia to ATP *in vitro* can augment their ramification index ^39^. Thus, we treated iMGLs with 1mM ATP for 6 h and measured its effect on their morphology. Indeed, we observed a significant increase in the ramification index of iMGLs upon ATP treatment (Supplementary Fig. S2G). Upon transduction of iMGLs with AAVs, we did not observe any significant changes in their ramification index at 24 h post-transduction when compared to the control condition (Supplementary Fig. S2H). However, at 72 h post transduction, we noted a decrease in the ramification index of AAV6 transduced iMGLs when compared to control and AAV8 transduced microglia (Fig. 6A-B). AAV6-transduced iMGLs also exhibited a hypertrophic morphology, which is associated with a microglial response to injury, infection or loss of homeostasis ^40^.

**Figure 6:**
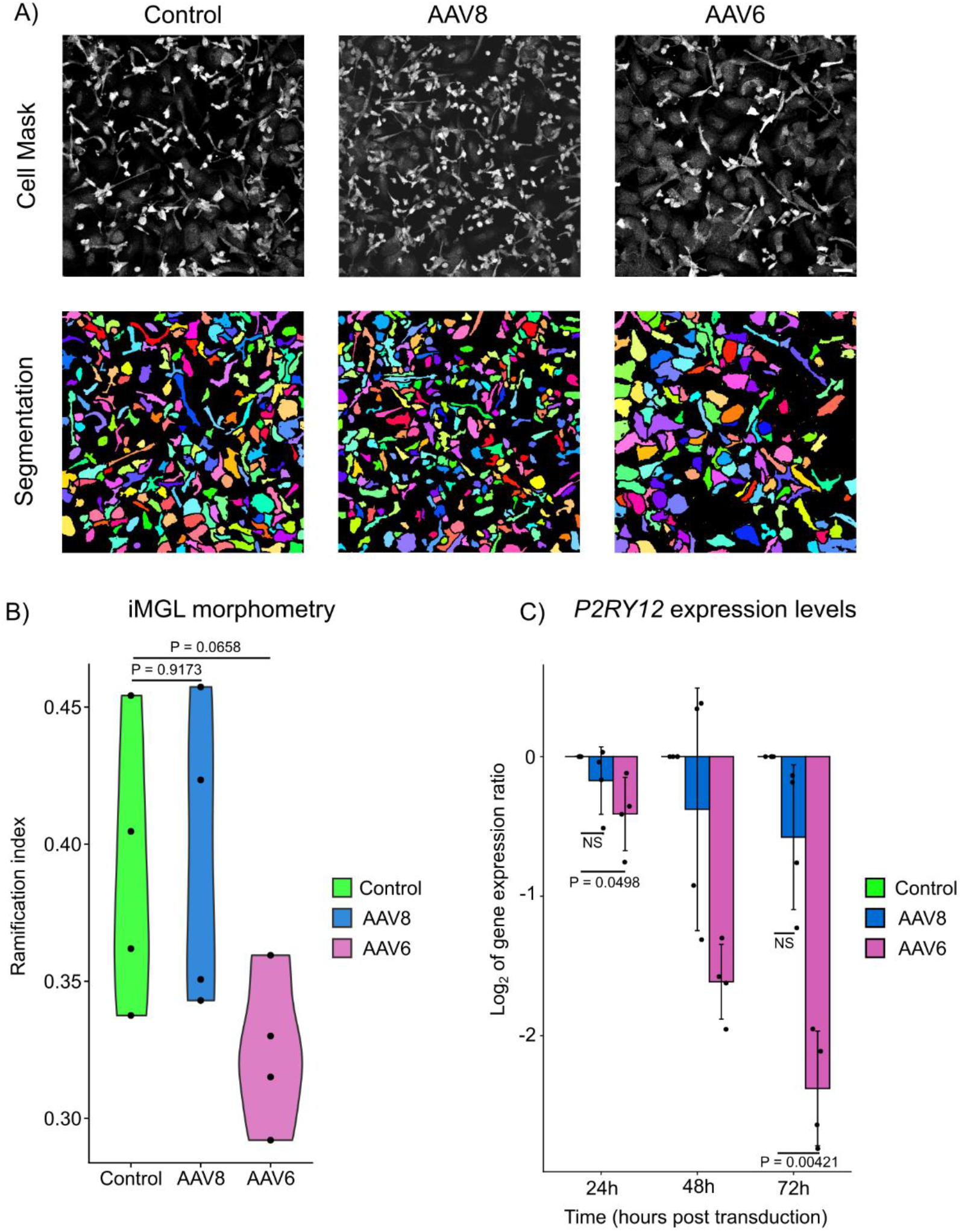
AAV6 transduction of iMGLs drives morphological changes and downregulates *P2RY12* expression levels. A) Representative micrographs of iMGLs depicting morphological changes as a function of AAV transduction. B) Violin plot depicting the ramification index of control iMGLs and iMGLs transduces with AAV8 and AAV6, respectively; each dot represents an individual biological replicate (n = 4 biological replicates). Statistical significance was calculated using a linear regression model. C) Histogram depicting the log2 value of gene expression ratio of *P2RY12* with respect to the housekeeping gene, *ACTB* at 24 h, 48 h and 72 h post transduction with AAV serotypes, AAV8 and AAV6, respectively. Data are plotted as mean ± standard deviation; each dot represents an individual biological replicate (n = 4 biological replicates). Statistical significance was calculated using the Kruskal-Wallis test with Dunn’s multiple comparison test. NS = not significant.

This prompted us to examine if the change in microglial morphology upon AAV6 transduction was accompanied by gene expression changes*. P2RY12* is a purigenic receptor, which plays an essential role in the tissue surveillance function of microglia and is a marker for homeostatic microglia. Notably, this gene is downregulated in microglia in the context of neuroinflammation ^41^. It has been shown that mouse microglia downregulate *P2ry12* upon AAV-mediated retinal inflammation ^13^. Upon LPS treatment, we detected a significant downregulation of *P2RY12* in iMGLs, confirming their suitability to model the microglial reactivity and transcriptional changes upon changes in the iMGL growth environment (Supplementary Fig. S2I). Next, we profiled the transcript levels of *P2RY12* upon AAV transduction at 24 h, 48 h and 72 h post-transduction. qPCR analysis revealed a gradual decrease in the levels of *P2RY12* expression over a period of 72 h in AAV6 transduced iMGLs (Fig. 6C). Moreover, this decrease was significant at 24 h and 72 h when compared to the control condition. In the case of AAV8-transduced iMGLs, while we observed a slight decrease in *P2RY12* expression levels, it was not significant when compared to the control condition.

Thus, morphological and molecular analysis revealed that AAV6 induced a change in the iMGL cell state, suggesting that iMGL reactivity upon AAV transduction is serotype-dependent.

### Elevated levels of intracellular AAV6 genomes triggers a nucleic acid sensing pathway in iMGLs

We wondered about the cellular mechanisms responsible for inducing microglial reactivity upon AAV6 transduction. We hypothesized if the levels of intracellular AAV genomes could be a determining factor for inducing microglial reactivity.

Thus, we employed RNAscope to assess the levels of intracellular AAV genomes at 24 h and 72 h post transduction. We used a DNA probe specific to the CAG promoter to detect intracellular viral genomes and a RNA probe for the *IBA1* transcript to identify the iMGL cell body. At 24 h post transduction, we observed a considerably higher number of fluorescent dots for the DNA probe in AAV6 transduced iMGLs as compared to AAV8 (Fig. 7A). We employed a semi-quantitative scoring method to assess if the number of intracellular AAV6 genomes was significantly higher than that of AAV8. By binning the cells into 4 distinct categories based on the number of DNA probe dots, we demonstrated that proportion of iMGLs with 15 or more CAG promoter probes per cell (Score 4) was significantly higher for AAV6 (92.43%) as compared to AAV8 (23.47%) (Fig. 7B). On the contrary, the proportion of iMGLs with 1-3 CAG promoter probes per cell was significantly higher for AAV8 (17.43%) as compared to AAV6 (0.0%) (Fig. 7B). To assess if the temporal dynamics of cellular entry differed between AAV6 and AAV8, we assessed the levels of intracellular viral genomes at 72 h post transduction.

**Figure 7:**
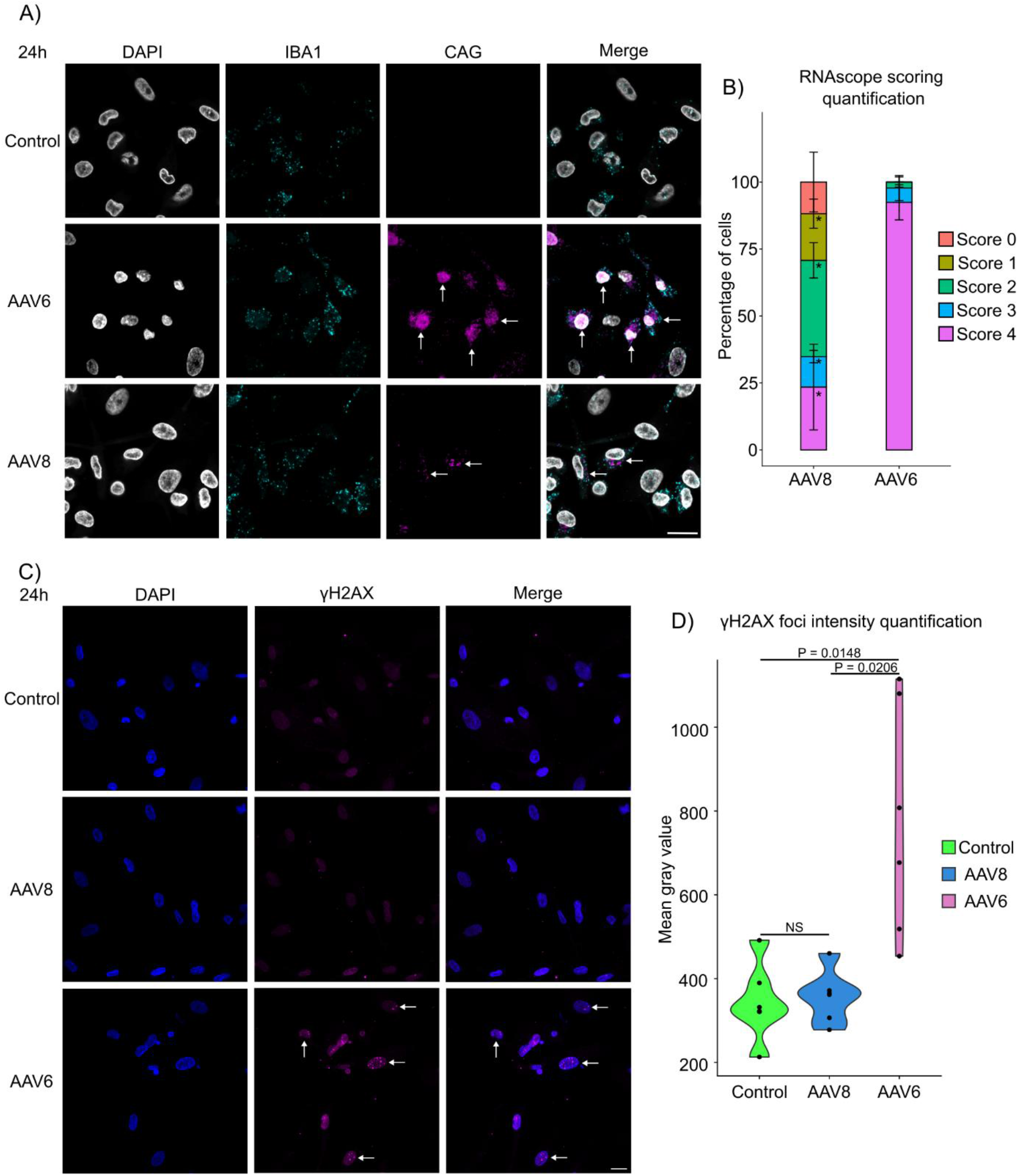
Elevated levels of intracellular AAV6 genomes triggers a DNA double strand break response in iMGLs. A) Representative micrographs of RNAscope staining of iMGLs transduced with AAV serotypes, AAV6 and AAV8, respectively at 24 h. The probe targeting the *IBA1* transcript is a RNA probe while the probe targeting the CAG promoter is a DNA probe. Filled arrowheads depict the transduced cells with intracellular AAV genomes. Scale bar 50 µm. n = 4 biological replicates). B) Stacked bar plot depicting the proportion of cells in each class of the ACD scoring system for RNAscope (n = 3 biological replicates). The asterisk ‘*’ denotes a statistically significant difference between the AAV6 and AA8 condition for Score 1, Score 2, Score 3 and Score 4, respectively. C) Representative micrographs of iMGLs stained with an anti-γH2AX antibody to visualize the presence of γH2AX foci upon AAV cellular entry. Filled arrowheads depict AAV6-transduced iMGLs with γH2AX foci. Scale bar = 20 µm. n = 6 biological replicates. D) Violin plot depicting the quantification of the mean gray value intensity of γH2AX foci in iMGLs under the given conditions. Each dot represents an individual biological replicate. n = 6 biological replicates. Statistical significance was calculated using the Kruskal-Wallis test with Dunn’s multiple comparison test. NS = not significant.

However, the number of viral DNA probe dots did not increase in AAV8 transduced iMGLs at this later time point, suggesting that AAV uptake by iMGLs is not dependent upon the time of exposure to the virus (Supplementary Fig. S4A). Thus, our results demonstrated that AAV6 is more efficiently uptaken by iMGLs than AAV8. Next, we hypothesized if the elevated intracellular levels of AAV6 genomes would trigger a nucleic acid sensing pathway, since the accumulation of fragmented and damaged DNA is a key feature of neurodegeneration-associated microglial reactivity ^42–44^. Thus, we decided to quantitatively assess the link between intracellular AAV genomes and induction of a DNA damage response in iMGLs. As a control, we treated iMGLs with etoposide, a topoisomerase II inhibitor, to induce DNA double-strand breaks and the subsequent appearance of γH2AX foci (Supplementary Fig. S4B). Furthermore, we measured the intensity of γH2AX foci using ImageJ, which increased significantly upon etoposide treatment when compared to the control condition (Supplementary Fig. S4B-C). Then, we asked if elevated levels of AAV6 genomes could trigger a DNA damage response in iMGLs. Indeed, we detected an increase in γH2AX foci in the AAV6 condition at 24 h post transduction when compared to control and AAV8-transduced iMGLs (Fig. 7C-D). Thus, we established a link between the accumulation of intracellular AAV genomes and the triggering of a DNA damage response in iMGLs, which is a key feature of neuroinflammation-associated microglial reactivity.

## DISCUSSION

In this study, we gained insights in the role of serotype dependent cellular uptake of AAV in driving IBA1+ cell reactivity and a potential mechanism by which the elevated levels of intracellular AAV vector genomes act as danger signal to drive microglial reactivity.

Upon subretinal delivery of AAV6, we observed significant bilateral ocular inflammation with diffuse retinal disorganization, migration of IBA1+ cells in outer retinal layer and an invasion of adaptive immune cells in the retina of 2 out of the 3 minipigs. Furthermore, the IBA1+ cells displayed morphological features associated with an inflammatory cell state ^10,34^. We also observed the infiltration of innate and adaptive immune cells, which is a hallmark feature of ocular and chorioretinal inflammation in various retinal disorders including diabetic retinopathy and AMD ^45^. Two factors pointed towards the risk of an inflammatory reaction following subretinal injection of AAV6-CAG-EGFP in minipigs. Firstly, AAV6 has been shown to induce an adaptive immune response with a single dose in rats, which is consistent with our observation of sequential bilateral ocular inflammation in 2 pigs injected with a single dose of AAV6 and with the infiltration of B lymphocytes in the retinas ^46^. Secondly, we used a high dose of vector (3E12 VG/ml in 150µL injection) to test the upper limits of vector doses used in clinical trial but which also increases the risk of a dose-dependent immune response ^47^. However, our data suggested that the inflammation was not due to the expression of EGFP in the IBA1+ cells but rather due to the transduction of IBA1+ cells by AAV6, since accumulation of fragmented double stranded DNA is a potent inducer of the type-I interferon signaling pathway and subsequent microglia-mediated inflammation ^12,14,48^. However, the assessment of such molecular mechanisms is experimentally challenging in large animal models. A major limitation for the *in vivo* part of this study is the limited number of animals, with only three minipigs injected with AAV6. Thus, the *in vivo study* is statistically underpowered to compare the inflammatory signs between the test and control animals.

To overcome these limitations, we employed iMGLs as an *in vitro* model to elucidate the cellular mechanisms driving microglial reactivity upon AAV transduction. Our phenotypic and molecular data *in vitro* demonstrated that AAV6 is a more efficient transducer of iMGLs when compared to AAV8. The elevated levels of intracellular AAV6 genomes also correlated with transition away from the homeostatic cell state, as evidenced by change in cell morphology and reduction in *P2RY12* expression levels. However, the change in iMGL cell state was not driven by supra-physiological levels of EGFP protein expression. Instead, our RNAscope data demonstrated that the level of intracellular AAV6 genomes in iMGLs *in vitro* was significantly higher when compared to AAV8 and control cells. This could be explained by the sensing of the vector genome by the various microglial pattern recognition receptors such as TLR9 ^49,50^. Furthermore, we made a mechanistic link between the sensing of elevated levels of intracellular AAV6 genomes and presence of cellular marker for DNA damage, which could lead to microglial reactivity ^21,43,44,51^. The results also highlights the significance of a nucleic acid sensing threshold mechanism since the low levels of intracellular AAV8 genomes did not trigger a similar response. This observation is in accordance with previous studies demonstrating the ability of AAV8 to transduce mononuclear phagocytic cells without activating them ^52^. The differential response of iMGLs towards AAV serotypes might be due to the influence of the AAV serotype on intracellular trafficking and future studies should be designed to elucidate this aspect of AAV-host cell interaction ^53^.

Our data underscores the need to select AAV serotypes or develop AAV vector variants that will either be de-targeted from microglia or efficiently transduce these innate immune cells without triggering an innate immune response. While an iMGL monoculture is suitable to model the cell autonomous response to AAV transduction, it can only partially recapitulate the complex AAV-mediated immune response observed *in vivo*. A recent study employing iMGLs to model AAV mediated inflammation attempted to overcome this limitation by transplanting the iMGLs into a chimeric mouse model ^54^. It has been demonstrated that the transplantation of human iMGLs into the mouse brain improve the morphological and molecular characteristics of these cells ^55^. However, the results of modeling the AAV mediated immune response using a chimeric model should be cautiously interpreted, given the species-specific differences of the immune system ^56^.

In summary, our study highlights the mechanism by which AAV uptake by microglial cells can lead to the gene therapy associated inflammation and will pave the way for the development of safe and efficacious AAV vectors for central nervous system gene therapy.

## Abbreviations

*AMD*: Age-related Macular Degeneration
*AAV*: Adeno-Associated Virus
*CNS*: Central Nervous System
*EGFP*: Enhanced Green Fluorescent Protein
*ERG*: Electroretinogram
*GCL*: Ganglion Cell Layer
*GTAU*: Gene-Therapy Associated Uveitis
*iPSC*: Induced Pluripotent Stem Cell
*iMGLs*: iPSC derived Microglia-like Cells
*IRD*: Inherited Retinal Disease
*IZ*: Injected Zone
*LPS*: Lipopolysaccharide
*NIZ*: Non-Injected Zone
*NHP*: Non-Human Primate
*OD*: Oculus Dexter
*OS*: Oculus Sinister
*ONL*: Outer Nuclear Layer
*OCT*: Optical Coherence Tomography
*RPE*: Retinal Pigment Epithelium

## DECLARATIONS

### Ethics approval

We confirm that all experimental protocols involving animals were approved by the Animal Experimentation Ethics Committee of Pays de la Loire (France) and the Ministry of Higher Education and Research (approval number: 36573-2022041310585859v3).

### Consent for publication

Not applicable

### Data availability statement

All data generated or analysed during this study are included in this published article (and its Supplementary Information files).

### Authors’ contributions

Study concept and design: JBD, AP, TC, OA. Obtaining funding: JBD, TC, OA. Acquisition, analysis of data: JBD, AP, JD, LL, ET, TC, BR, AM, DA, VB, TL, AL, LC, PS, MG, MJ, CC. Interpretation of data; JBD, AP, JD, LL, ET, MG, CC, MJ, CLG, OA, TC, GL. Supervision of JBD: TC, OA, MW, GL. Drafting of the manuscript: JBD, AP OA and TC. Approval of the submitted version of the manuscript: all authors.

## Acknowledgements

We thank the staff of the vector core ViVeM (Viral Vector Manufacturing Center) of TaRGet laboratory (Nantes, awarded IBB France 2030) for the production of all AAV vector batches and Nicolas Jaulin for this advice and assistance on the FACS experiments. We also thank the staff of the Boisbonne Centre (ONIRIS, Nantes-Atlantic College of Veterinary Medicine, Nantes) where animal research took place. We acknowledge Mélodie Ambroset and Philippe Hulin from the MicroPICell core facility (SFR Bonamy, BioCore, Inserm UMS 016, CNRS UAR 3556, Nantes, France), member of the Scientific Interest Group (GIS) Biogenouest, IBISA, and the national infrastructure France-Bioimaging supported by the French national research agency (ANR-24-INBS-0005 FBI BIOGEN) for their advice assistance with confocal microscopy and image analysis. The authors acknowledge the Cytocell - Flow Cytometry and FACS core facilty (SFR Bonamy, BioCore, Inserm UMS 016, CNRS UAR 3556, Nantes, France) for its technical expertise and help, member of the Scientific Interest Group (GIS) Biogenouest and the Labex IGO program supported by the French National Research Agency (n°ANR-11-LABX-0016-01). The authors acknowledge THERASSAY core facility (SFR Bonamy, l’Institut du Thorax UMR 1087, CNRS UMR 6291, Nantes, France) member of the Scientific Interest Group (GIS) Biogenouest and IBISA, for the use of the Bioplex 200 and their technical support.

## Supplementary figure legends

**Supplementary Figure 1:**
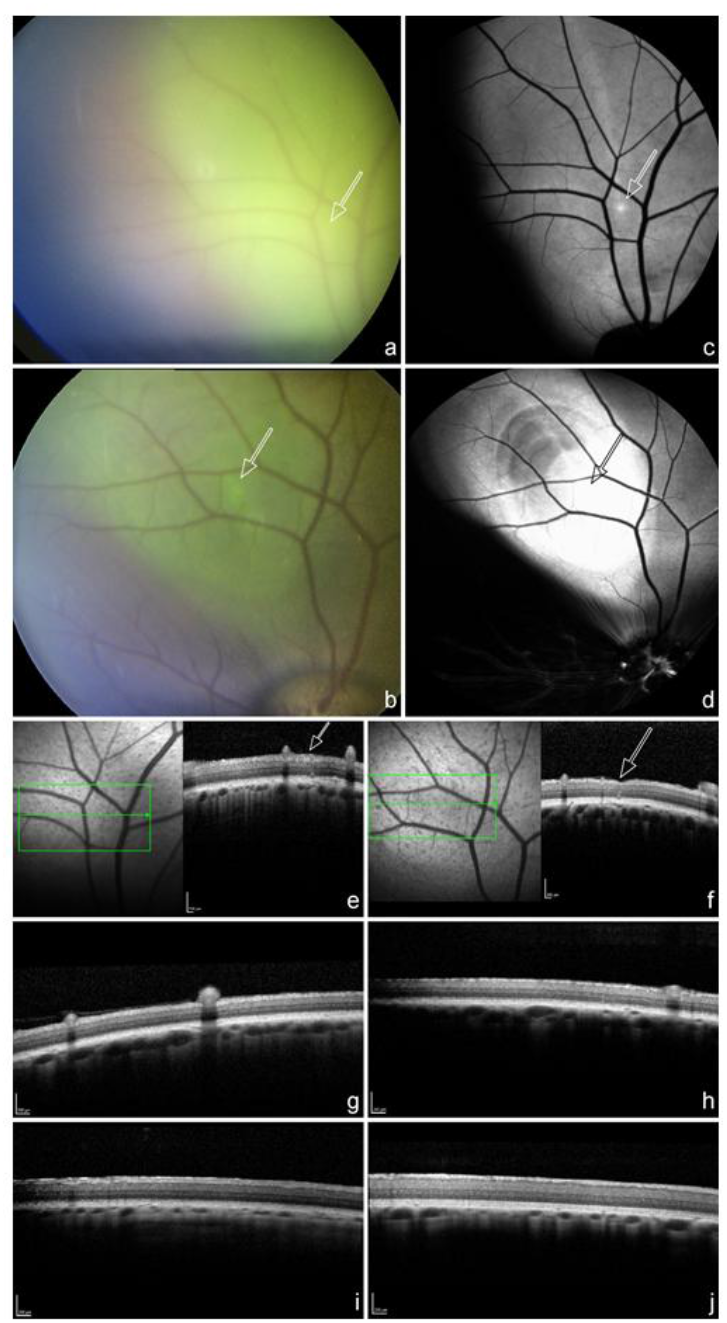
Color fundus and OCT analysis. Color fundus images show the EGFP expressing region (a, b) corresponding to autofluorescence fundus images of AAV injected retinas (c, d). The retinal cross-section analysed by OCT at the site of injection (e, f and i, j) is compared to peripheral regions (g, h).

**Supplementary Figure 2:**
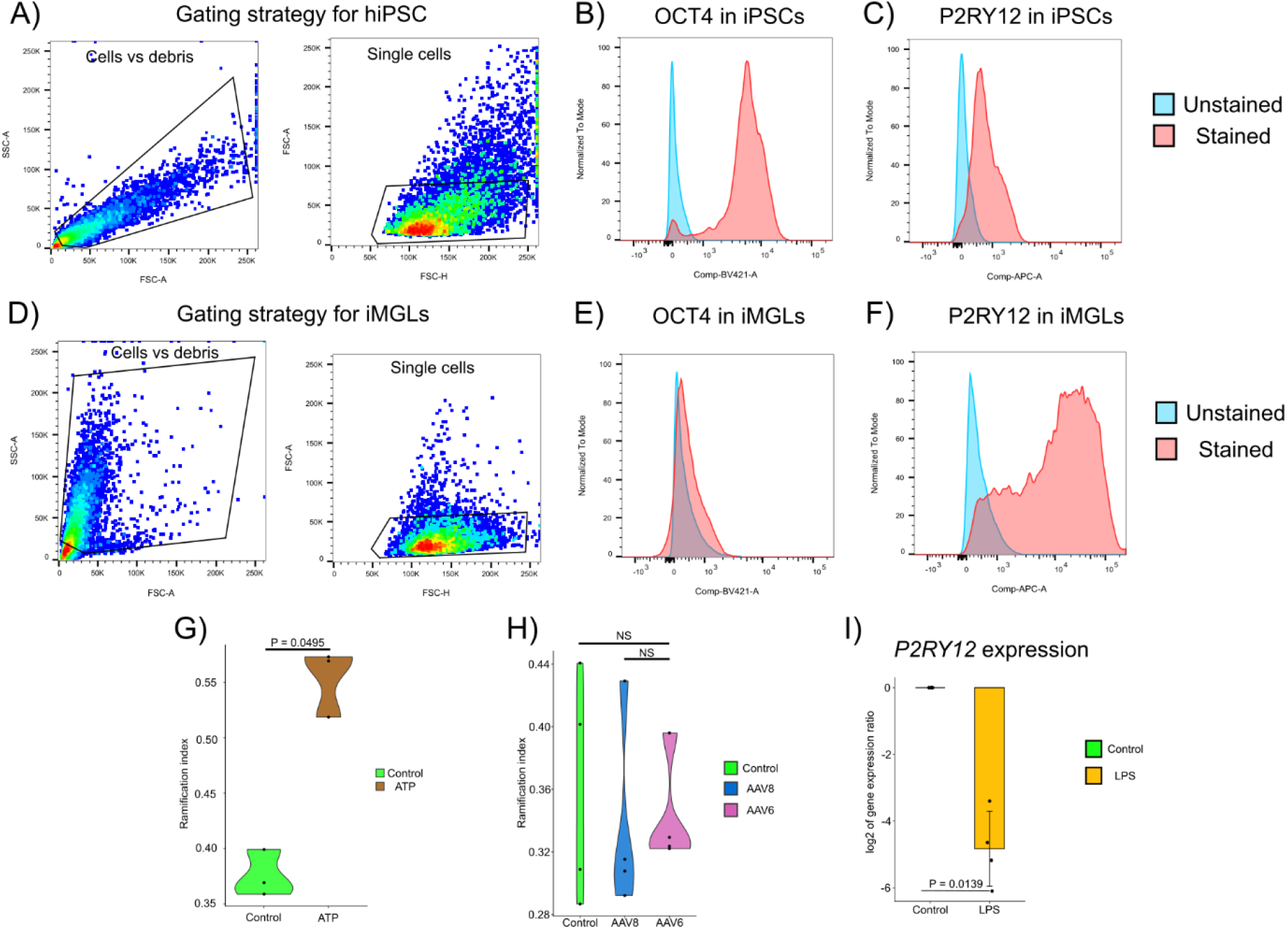
iMGL differentiation quality control. A) Flow cytometry gating strategy for the iPSC lines used in the study. B-C) Plots depicting the protein expression levels on OCT4 and P2RY12 in iPSCs, respectively. D) Flow cytometry gating strategy for the iMGLs used in the study. E-F) Plots depicting the protein expression levels on OCT4 and P2RY12 in iMGLs respectively. G) Violin plot depicting the ramification index of iMGLs upon ATP treatment for 6 h (n = 3 biological replicates) H) Violin plot depicting the ramification index of control iMGLs and iMGLs transduced AAV8, and AAV6, respectively at 24 h post transduction (n = 4 biological replicates). I) Histograms depicting the log_2_ value of gene expression ratio of *P2RY12* with respect to the housekeeping gene, *ACTB* at 24 h post LPS treatment (n = 4 biological replicates). Statistical significance was calculated using the Kruskal-Wallis test with Dunn’s multiple comparison test. NS = not significant.

**Supplementary Figure 3:**
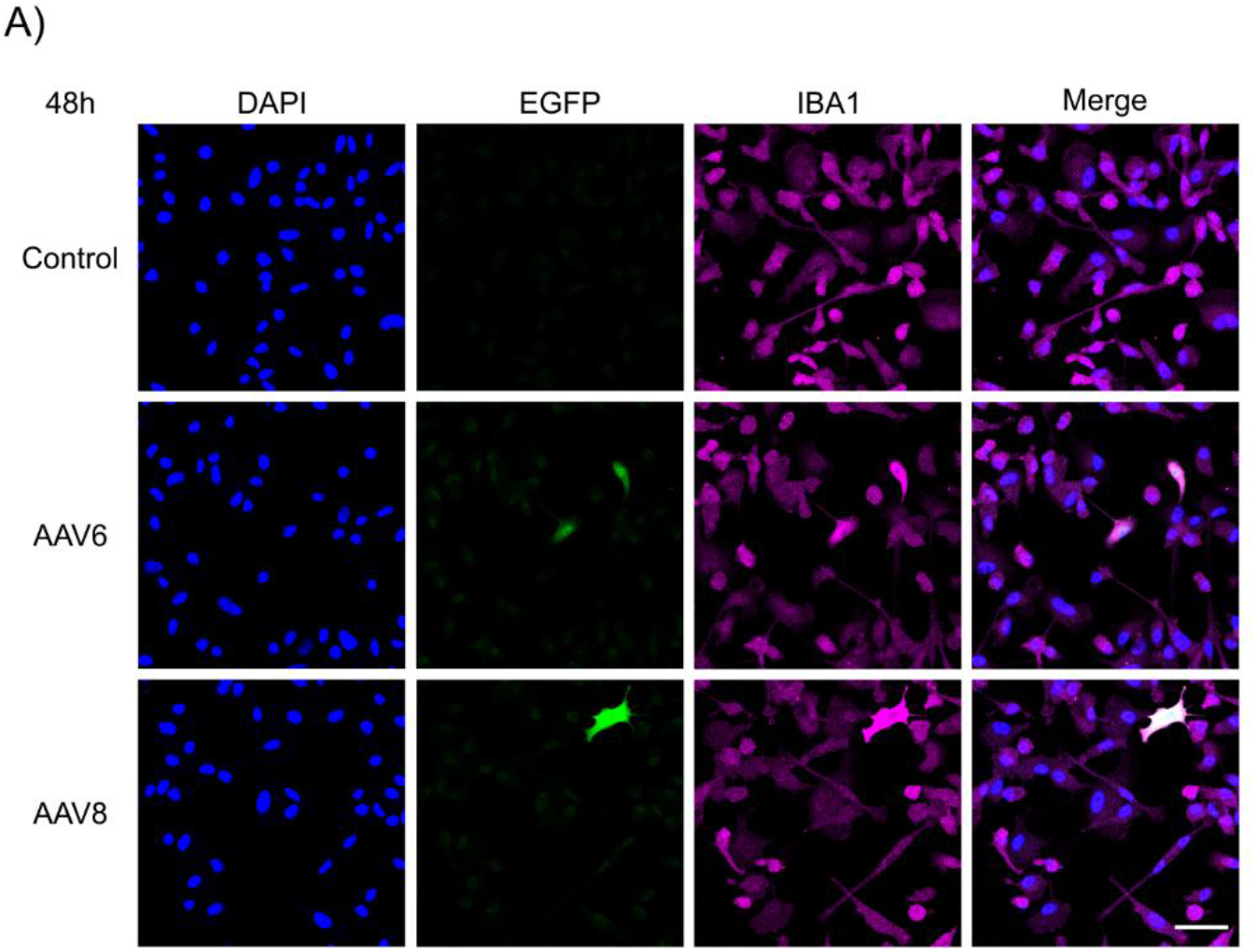
iMGLs are not permissive to AAV-mediated transgene expression at 48 h post transduction. A) Representative micrographs of iMGLs transduced with AAV serotypes, AAV6, and AAV8, respectively at 48 h post transduction. Scale bar 50 µm. n = 4 biological replicates.

**Supplementary Figure 4:**
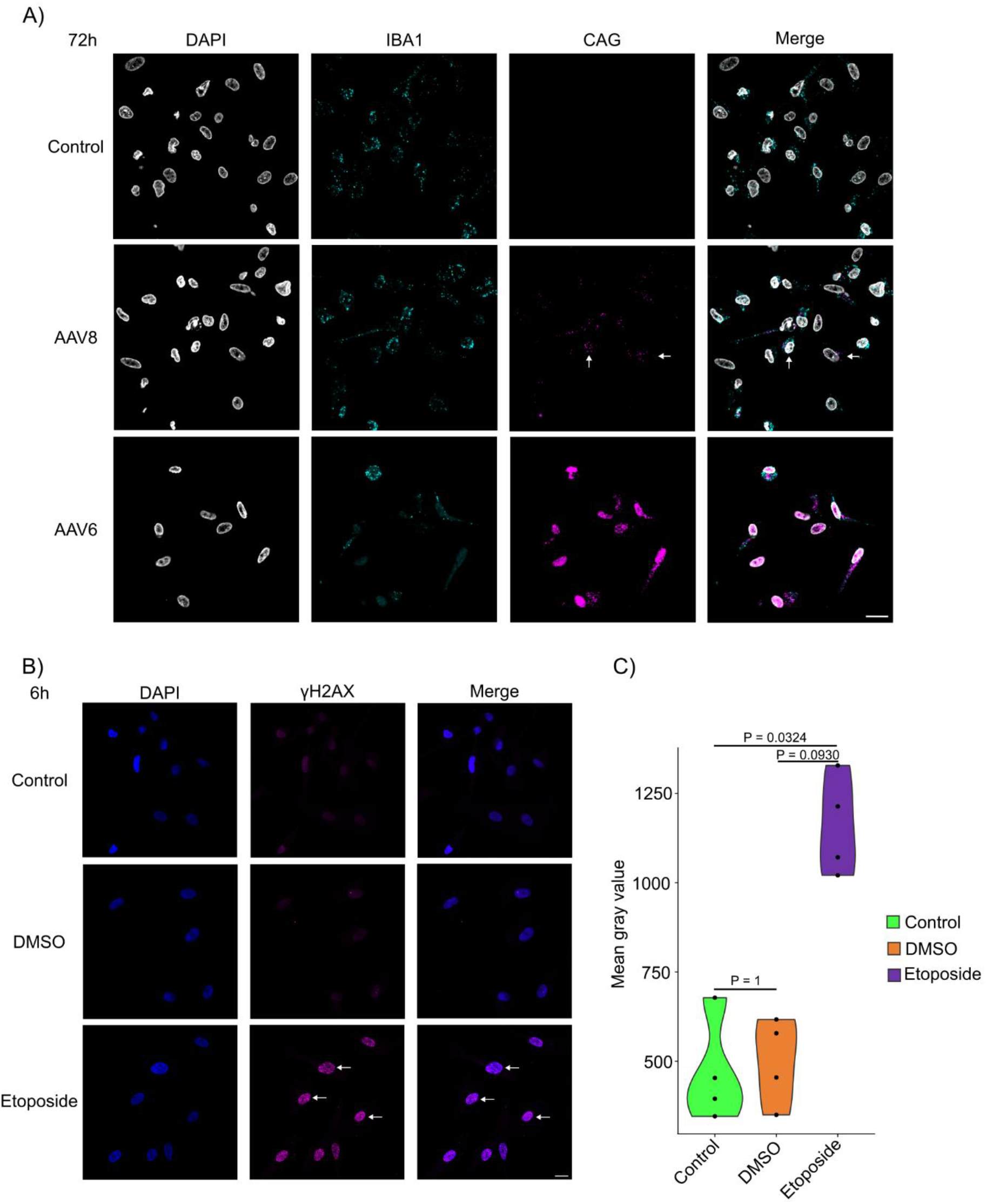
Transduction of iMGLs by AAV vectors is serotype dependent. A) Representative micrographs of RNAscope staining of iMGLs transduced with AAV serotypes, AAV6, and AAV8, respectively at 72 h post transduction. The probe targeting the *IBA1* transcript is a RNA probe while the probe targeting the CAG promoter is a DNA probe. Filled arrowheads depict the transduced cells in the condition AAV8. Scale bar 50 µm. n = 2 biological replicates. B) Representative micrographs of iMGLs stained with an anti-γH2AX antibody to visualize the presence of γH2AX foci upon treatment with DMSO (vehicle control) and 50 µM etoposide. Filled arrowheads depict etoposide-treated iMGLs with γH2AX foci. Scale bar = 20 µm. n = 4 biological replicates. C) Violin plot depicting the quantification of the mean gray value intensity of γH2AX foci in iMGLs under the given conditions. Each dot represents an individual biological replicate. n = 4 biological replicates. Statistical significance was calculated using the Kruskal-Wallis test with Dunn’s multiple comparison test. NS = not significant.

## Supplementary table legend

**Supplementary table 1:** Number of pig eyes used in this study. ERG: electroretinogram; OCT: optical coherence tomography; IHC; Immunohistochemistry; IHC and RNAscope were performed on retinal section. * *In vivo* imaging was not possible due to inflammation in two out of the 3 AAV6-injected pigs.

| Number of pig eyes used in this study |  |  |  |  |  |  |
| --- | --- | --- | --- | --- | --- | --- |
| Treatment | *In vivo* analysis |  |  | Post-mortem analysis (4 weeks p.i) |  |  |
|  | Fundus | OCT | ERG | IHC |  | RNAscope |
|  |  |  |  | Flat-mount | Retinal section |  |
| Uninjected | 2 | 2 | 2 | 2 | 0 | 0 |
| Saline | 4 | 0 | 4 | 2 | 2 | 2 |
| AAV6 | 2* | 2* | 4 | 3 | 3 | 2 |
| AAV8 | 2 | 2 | 0 | 2 | 2 | 2 |

**Supplementary table 2:** List of antibodies used in this study.

| Type | Name | Host | Dilutiion | Reference |
| --- | --- | --- | --- | --- |
| Primary antibodies | IBA1 | Rabbit | 1 in 400 | PA527436 |
|  | CD3 | Mouse | 1 in 200 | 916MSM10-P0 |
|  | CD79 | Rabbit | 1 in 100 | 973-RBM10-P0 |
|  | CD8 | Mouse | 1 in 400 | MA5-28520 |
|  | CD4 | Mouse | 1 in200 | MCA1749GA |
|  | CD14 | Mouse | 1 in 400 | MCA1218GA |
|  | CD163 | Mouse | 1 in 100 | MCA2311GA |
|  | IBA1 | Goat | 1 in 500 | ab5076 |
|  | GFP | Chicken | 1 in 300 | ab13970 |
|  | p-histone H2A.X | mouse | 1 in 200 | 05-636-l |
|  | OCT4, BV421 |  | 1 in 100 | 653712 |
|  | P2RY12, APC |  | 1 in 100 | RRID: AB_2910451 |
| Secondary antibodies | Rabbit | Goat | 1 in 500 | A11010 |
|  | Mouse IgG1 | Goat | 1 in 1000 | A21127 |
|  | Rabbit IgG1 | Donkey | 1 in 1000 | A31573 |
|  | Mouse IgG2a | Goat | 1 in 1000 | A21241 |
|  | Mouse IgG2b | Goat | 1 in 1000 | A21242 |
| Lectine PNA |  |  | 1 in 250 | L34260 |

## Materials and Methods

All experiments were performed under the control of our quality management system, which has been approved by Lloyd’s Register Quality Assurance to meet the requirements of the international management system standard ISO 9001:2015. This covers a range of laboratory activities, including research experiments and the production of research-grade viral vectors.

### Animal Care and welfare

Experiments were performed on six female Yucatan Mini-pig provided by INRAE (Saint Gilles, France). They were 6 months old and weighed around 20kg. The experiments were conducted at the Boisbonne Centre (ONIRIS, Nantes-Atlantic College of Veterinary Medicine, Nantes). All animals were handled in accordance with the ARVO Statement for the Use of Animals in Ophthalmic and Vision Research. The experiments involving animals were conducted in accordance with agreement delivered by the Animal Experimentation Ethics Committee of Pays de Loire (France) and the Ministry of Higher Education and Research (authorization APAFIS.36573-2022041310585859v3).

### Vectors

All AAV-CAG-EGFP vectors were produced by the Viral Vector Manufacturing Center (ViVEM) in Nantes using HEK293 adherent cells and calcium-based transfection method. Vectors were purified by employing a cesium-chloride gradient. Titers were determined in viral genomes/ml using a quantitative PCR method. The use of AAV vectros were approved by the Scientific Committee of the “Haut Conseil des Biotechnologies” of the French Ministry of Research.

### Surgery

Pupils were dilated with atropine 0.3%, tropicamide 0.5% and neosynephrine 10% eye drops. General anesthesia was induced by intramuscular injection of tilétamine / zolazépam (Zoletil ®, 4 mg/kg) and atropine (0,05mg/kg) and maintained with isoflurane (1-2 %) / O2 (1-1.5L/min). Analgesia was performed with pre-operative intravenous buprenorphine (0,01 mg/kg) and then repeated every 6 hours. Local anesthesia was associated with oxybuprocaïne 0.4% eye drops. Ocular surface was disinfected by povidone iodine 5%. After lateral canthotomy, a 25-gauge three port vitrectomy was performed with posterior vitreous detachment. Vitreous sampling was performed with the vitrector, at the very beginning of the core vitrectomy, before the activation of the saline solution infusion. Bleb retinal detachment was performed by subretinal injection of 150µL of viral vector or saline solution (BSS) along the temporal superior vascular arcade with a 41-gauge subretinal injection cannula. Sclerectomies and lateral canthotomy were sutured with vicryl® 7/0 (adsorbable). To replicate the conventional clinical preventive anti-inflammatory treatment, subconjunctival injection of 0.5ml of long-acting corticosteroid (methylprednisolone, 40mg/ml) was performed at the end of the procedure in each eye. Ophthalmic antibiotic ointment (chloramphenicol) was administered on the eye at the end of the procedure and for two days afterwards.

### Examinations

All the following *in vivo* examinations were performed at least one week before surgery and one month after surgery (Supplementary table 1). Blood (2 mL) was collected from a vein of the ear. Pupils were dilated with tropicamide 0.5% and neosynephrine 10% eye drops. General anesthesia was induced by intramuscular injection of tilétamine / zolazépam (Zoletil®, 4 mg/kg) and atropine (0,05mg/kg) and maintained with isoflurane (1-2 %) / O2 (1-1.5L/min). Color fundus photographs were taken with a Canon© CF-60 UVi retinal camera. Eye fundus and Optical coherence tomography (OCT), were performed with the Spectralis Heidelberg© Retinal Angiograph (HRA). OCT B-scans were performed on the whole macular area. Total retinal thickness (RT) and outer nuclear layer thickness (ONL) were measured on optical coherence tomography (OCT) of the injected eyes for the AAV8 and AAV6 condition before injection and 1 month after injection, in 3 random points in the IZ and in the macular NIZ.

### Electroretinogram (ERG)

Pupils were dilated with atropine 0.3%, tropicamide 0.5% and neosynephrine 10% eye drops. General anesthesia was induced by intramuscular injection of tilétamine / zolazépam (Zoletil®, 4 mg/kg) and atropine (0,05mg/kg) and maintained with isoflurane (1-2 %) / O2 (1-1.5L/min). Local anesthesia was associated with oxybuprocaïne 0.4% eye drops. Pigs were dark-adapted for 20 minutes. Carbomère (Gel Larmes, Thea, France) was applied to each eye to prevent corneal dehydration during recordings. A computer-based system MonColor (Metrovision, Pérenchies, France) and contact lens electrodes (ERG-jet; Microcomponents SA, Grenchen, Switzerland) were used. ERG max were recorded in a standardized manner according to the International Society for Clinical Electrophysiology of Vision (ISCEV), with flashs 3.0 cd.s/m² at 10 seconds intervals.

### Flat mounted retinal examination

Immediately after the last eye examinations, pigs were euthanized with intravenous high dose of phenobarbital (180mg/kg) and carefully enucleated, following confirmation of death by a veterinarian. Retinas of the right eyes were flat mounted and stained with an IBA1 antibody and DAPI (Supplementary table 2). Imaging was performed with a Nikon A1R confocal microscope (Champigny/Marne, France) with NIS v 5.30.03 using an x25 Plan Apo objective silicone immersion N.A. 1.05. Round and well-defined IBA1+ cells were counted on confocal slices of flat-mount retinas from the right eyes of the pigs in 6 random squares of 500 µm side length in the injected zone (IZ) and in non-injected zones (NIZ). If a vessel was present in the square, cells visible in the vessel and near the vessel of at most 1.5 vessel diameter were not counted. For all squares, an image stack was acquired. Each stack acquisition was analyzed and each IBA1+ cell were contoured under NIS and the manual measurement function. Manual threshold was applied for each cell on a stack projection and NIS calculates for each cell ShapeFactor (4 x pi x aire) / perimeter²).

### Immunohistochemistry

Retinas were cryosectioned (12-µm sections). Tissue sections were fixed in PFA 4% for 10 min at RT and then washed 3 times with 1x PBS for 5 min. Sections were permeabilized with PBS 0.2% Triton for 10 min at room temperature (RT), washed 3 times with 1x PBS for 5 min and blocked for 45 min at RT in blocking solution (5% bovine serum albumin, 2% goat serum, 10% primate serum, in 1X PBS). For immunostaining, primary antibodies were diluted (Supplementary table 2) in blocking solution and incubated overnight at 4°C. In the negative control condition, no primary antibody was added. The sections then were washed 3 times with 1x PBS for 10 min at RT and incubated with appropriate secondary antibodies (Supplementary table 2) in blocking solution for 1 hour at RT. The nuclei were counterstained with 4,6-diamidino-2-phenylindole (DAPI) diluted with 1x PBS 1 :1000 for 20 min in the dark at RT. Finally, sections were washed 3 times with 1x PBS for 10 min at RT and mounted under a glass coverslip using Prolong Gold Antifade Mountant.

### RNAscope *in situ* hybridization for *in vivo* retinal sections and *in vitro* iMGL cells

Frozen Yucatan minipigs retina tissues were cryosectioned at -20°C to 12-μm thickness and mounted on glass slides. Serial sections were collected, and slides were dried at -20°C for 1 hour before storage at -80°C until use. RNA *in situ* hybridization for *IBA1*, *EGFP*, and DNA hybridization for CAG were performed using RNAscope Multiplex Fluorescent Reagent Kit v2 Assay (ACDbio) according to the manufacturer’s instructions. Slides were removed from -80°C freezer and placed immediately into cold (4°C) 4% PFA, then incubated at 4°C for 1 hour. Slides were dehydrated in graded ethanol series (50%, 70%, 100% x2, 5 mins in each, at RT) then air-dried 5 mins at RT. Sections were ringed using hydrophobic barrier pen (ImmEdge, Vector Labs) and then treated with RNAscope hydrogen peroxide (ACDbio) for 10 mins at RT. After washing twice in distilled water, slides were protease treated (RNAscope Protease IV, ACDbio) for 30 mins at RT and washed a further two times in distilled water before use in RNAscope multiplex assay. Probe mixes used were as follows:

– RNAscope 3-plex positive control probe: POLR2A (318811-C1), PPIB (428591-C2) and UBC (400641-C3)
– RNAscope 3-plex negative control probe set (320871): Bacterial DapB (in C1, C2 and C3)
– RNAscope pig’s retina probe set: CAG-promoter-03 (1240811-C1), Ss-AIF1 (1110621-C2) and EGFP-04 (538851-C3)

Fluorophores were assigned as follows: probe (C1), TSA Vivid 650 (323273); probe (C2), TSA Vivid 570 (323272); 3’ probe (C3), TSA Vivid 520 (323271).

The remainder of the protocol was completed, including probe, Multiplex FL v2 AMP and HRP, and TSA Vivid Fluorophore incubations, all at 40 °C using the ACD HybEZ II Hybridation System. Washes were completed with RNAscope wash buffer. After RNAscope labelling, nuclei were stained with DAPI (ACDbio) for 30 sec, and slides were mounted in Prolong Gold Antifade mounting medium (Thermo Fisher) and allowed to dry overnight at RT. Images were acquired on a Nikon A1R confocal microscope.

RNAscope was performed on iMGLs cultured in ibiTreat µ-Slide 8 Well chamber slides (80826, Ibidi) with the following modifications. Briefly, 15,000 cells were seeded onto each µ well respectively. After 14 days of differentiation, the cells were fixed using 4% PFA (sc-281692, Santa Cruz Biotechnology) for 10 minutes at RT. Upon fixation and dehydration, the chamber slides were stored at 4^0^C. The cells were rehydrated in graded ethanol series (70%, 50% x2, 2 minutes in each) followed by 1x PBS for 10 minutes at room temperature. The cells were treated with Protease III for 10 minutes at room temperature. The cells were incubated with the probe C1/3 for 30 minutes at 40°C, followed by an incubation with RNAscope multiplex FL v2 HRP blocker for 15 minutes 40°C. The cells were incubated with RNAscope multiplex FL v2 Amp1/3 for 30 minutes at 40°C. The incubation with the TSA plus fluorophores was performed at 40°C for 30 minutes. The cells were stained with DAPI for 10 minutes at room temperature.

### Cytokine secretion assay

Cytokine levels in the anterior chamber fluid were quantified using a Luminex-based multiplex assay. Samples (50 µl) were collected before and after the sub-retinal injection and were analyzed using the MILLIPLEX Human Cyrokine/Chemokine/Growth Factor Panel A magnetic beads panel (PCYTMG-23K-19PX, Merck Millipore), according to the manufacturer’s instructions. Briefly, standards, controls, and samples were incubated with the antibody-immobilized beads. After washing, detection antibodies and Steptavidin-Phycoerythrin were added sequentially. The plates were read using a Bioplex 200 Luminex analyzer, and the data were processed using the Bio-Plex Manager software. Median Fluorescence Intensity (MFI) was used to calculate the cytokine concentrations based on a 5-parameter logistic curve-fitting method.

### iPSC culture

Two commercially available iPSC lines, SCTi003-A (Stemcell Technologies) and SK002.1 (Reprocell) were employed in this study. Both the iPSC lines have been ethically sourced and collected using IRB protocols by their respective commercial suppliers. The iPSC lines were routinely screened and tested negative for mycoplasma infection by quantitative PCR. The working cell banks of both hiPSC lines were confirmed to be karyotypically stable.

Briefly, iPSC were cultured in mTESR plus medium (100-0276, Stemcell Technologies) on matrigel (Corning) coated tissues culture grade dishes. There were passaged at 70% confluency using solution versene (15040066, Gibco). Cells were incubated at 37°C and 5% CO_2_ and fed every two days with fresh mTESR plus medium.

### Generation of iPSC derived microlia-like cells (iMGLs)

#### Embryoid body formation

iMGLs were obtained using a previously published protocol ^35^. Briefly, an Aggrewell 800 plate (34815, Stemcell Technologies) was employed for the formation of embryoid bodies (EBs). 0.5 ml of anti-adherence solution (07010, Stemcell Technologies) was added to each well. The plate was centrifuged at 3000 x g for 3 minutes. The anti-adherence solution was discarded by aspiration and the well was washed once using 1ml of 1x PBS. The PBS was discarded by aspiration and 1ml of EB medium containing mTESR plus, 50 ng/ml BMP-4 (AF-120-05ET, Peprotech), 50 ng/ml VEGF (100-20, Peprotech), 20 ng/ml SCF (300-07, Peprotech) and supplemented with 10 µM ROCK inhibitor (3746170, Sigma Aldrich) was added to each well. The Aggrewell 800 plate containing the medium was placed back into the incubator, during the preparation of the iPSCs. The iPSCs were assessed to identify over-proliferative colonies and spontaneously differentiating cells, which were eliminated by aspiration. The cells were washed once with 1x PBS and dissociated using solution Versene. The iPSCs were incubated in solution Versene at room temperature for 3-5 minutes. The cells were further dissociated using a cell scrapper. The cells were counted using a Countess 3 automated cell counter. The cell density per well were as follows: for SCTi003-A – 900,000 cells/ml and for SK002.1 – 700,000 cells/ml The cell suspension containing the required number of cells was centrifuged at 400 x g for 5 minutes. The cell pellet was resuspended in EB medium + ROCK inhibitor and 1ml of the cell suspension was added to the Aggrewell (total volume in the well 2ml). The Aggrewell 800 plate was centrifuged at 100 x g for 3 minutes. The embryoid bodies in the Aggrewell 800 plates were fed with EB medium. Briefly, 1ml of medium was aspirated from the top of the Aggrewell using a P1000 pipette tip and replenished with 1ml of fresh EB medium. This step was repeated to give a 75% medium change overall. The embryoid bodies were fed for up to 3 days in the Aggrewell.

#### Setting up of macrophage precursor factory

On day 4, the embryoid bodies were harvested as follows: The serological pipettes and falcon tubes used for the harvest of the EBs were coated with 1% BSA solution to prevent the adherence of the EBs to the surface of the pipettes and tubes. Using a 5ml serological pipette, the EBs were gently dislodged from the microwells and transferred onto an inverted 40 µm cell strainer (Corning, 352340), which was placed on top of a 50 ml falcon tube. The dislodging step of the EBs was repeated once to ensure maximum recovery. The cell strainer containing the EBs was turned over onto a new 50 ml falcon tube and the EBs were washed into the tube using 5ml of factory medium containing Advanced DMEM/F12 (Fisher Scientific, 12634028), 1x glutamax (100x, Thermofisher Scientific, 35050028), 15 mM HEPES solution (Thermofisher Scientific, 15630080), 5 µg/ml human recombinant insulin solution (Thermofisher Scientific, RP-10908), 15 µM tropolone (Sigma Aldrich, T-89702-1G), 25 ng/ml IL-3 (Peprotech, 200-03), 100 ng/ml M-CSF (Peprotech, 300-25), 100 U/ml Penicillin-Streptomycin (Thermofisher Scientific, 15070063). The EBs were seeded onto matrigel coated T-75 flasks and incubated at 37°C and 5% CO_2_ for 1 week with disturbance. Thereafter, the T-75 flasks were fed once per week with factory medium (50% of the original volume of the flask). This was repeated for 3-4 weeks until macrophage/microglia precursors appeared in the supernatant.

#### Differentiation of iMGLs

The macrophage/microglia precursors were harvested and differentiated to iMGLs as follows: The supernatant was gently harvested from T-75 factory flasks.The T-75 factory flask was replenished with 50% of conditioned factory medium and 50% of fresh factory medium. The harvested supernatant was passed through a 40 µm cell strainer to remove any clumps of cells or floating EBs. The filtered cell suspension was centrifuged at 400 x g for 5 minutes. The resulting cell pellet was resuspended in iMGL medium containing Advanced DMEM/F12 (Fisher Scientific, 12634028), 1x glutamax (100x, Thermofisher Scientific, 35050028), 50 U/ml Penicillin-Streptomycin (Thermofisher Scientific, 15070063), 100 ng/ml IL-34 (Peprotech, 200-34), 50 ng/ml TGF-β1 (Peprotech, 100-21C), 25 ng/ml M-CSF (Peprotech, 300-25), 10 ng/ml GM-CSF (Peprotech, 300-03) at the desired cell density. A 50% medium change was performed every 3-4 days. The iMGLs were used for experiments after 2 weeks of differentiation.

#### Fluorescence-activated cell sorting

Cells were harvested by washing them once with 1x PBS followed by their detachment using solution versene (Thermofisher Scientific, 15040066) at 37°C and 5% CO_2_ for 10 minutes. A cell scrapper was detach the cells and the resulting cell suspension was centrifuged at 400 x g for 5 minutes. The cell pellet was fixed using 2% paraformaldehyde on ice for 20 minutes. The fixed cells were incubated with the optimal primary antibody solution for 20 minutes on ice (see Supplementary table 2 for the primary antibody dilutions). The stained cells were washed once with 1x PBS, centrifuged at 400 x g for 5 minutes and resuspended in 1x PBS for the FACS analysis.

To stain for intracellular proteins, the cells were fixed and permeabilised using Cyto-Fast^TM^ Fix/Perm solution (426803, Biolegend) for 20 minutes at room temperature. The fixed and permeabilised cells were washed using 1x Cyto-Fast^TM^ Perm Wash solution and incubated with the optimal antibody solution prepared in 1x Cyto-Fast^TM^ Perm Wash for 20 minutes at room temperature. The stained cells were washed once with 1x Cyto-Fast^TM^ Perm Wash. The cells were centrifuged at 400 x g for 5 minutes, the cell pellet was resuspended in Cell staining buffer before the FACS acquisition. FACS was performed using a FACS Aria III (BS Biosciences). The gating the strategy was set by forward, side scatter and using isotype-control stained cells as the negative control.

#### Immunocytochemistry

Immunostaining of iMGLs was performed in ibiTreat µ-Slide 8 Well chamber slides (80826, Ibidi). Briefly, 15,000 cells were seeded onto each µ well respectively. After 14 days of differentiation, the cells were fixed using 4% paraformaldehyde for 10 minutes at room temperature. The fixed cells were washed twice with 1x PBS. The cells were incubated in blocking solution (3% BSA, 0.5% Triton X-100) at room temperature for 1 hour. The cells were incubated with the optimal primary antibody dilution in blocking solution over-night at 4°C. The following antibodies were used: Goat anti IBA1 (Abcam, ab5076), Chicken polyclonal anti-GFP (Abcam, ab13970). After washing the cells thrice with 1x PBS, the cells were incubated with the secondary antibody solution at room temperature for 1 hour. After washing the cells thrice with 1x PBS, the cells were stained with DAPI (Sigma Aldrich, D9542, 1:1000 diltion in blocking solution for 15 minutes at room temperature. Images were acquired using a Nikon A1R confocal microscope.

#### iMGL morphometry

The morphometric analysis of iMGLs upon AAV transduction was performed by employing Microglia Morphometry, a Fiji plugin, as described previously ^37^. Briefly, single frame confocal microscopic images were acquired for iMGLs infected with AAV6 or AAV8 at an MOI of 10^5^ for 24, 48 and 72 hours, respectively. The morphometric measurements of the control and AAV-transduced iMGLs were performed in a blinded manner. The plugin, ‘New Microglia Segmentation and Tracking’ was employed to semi-automatically segment the cells. The segmentation parameter, *minimal cell size*, was set to 150 µm^2^. Image smoothing was performed using an anisotrophic diffusion filter, followed by binarisation and conversion to a label mask for each cell. The generated label masks were manually corrected using the ‘Erase’ and ‘Draw’ tools in Fiji. A second plugin, Measure Microglia Morphometry’, was employed to obtain 17 morphometric measures. The ramification or branchness of the iMGLs was calculated using the following formula: Branchness = SkeletionNumBranchPoints / GeodesicDiameter_Pixel. The ramification index was calculated using the following formula: Ramification index = 2*Pi*Branchness.

#### RNAscope quantification

The RNAscope images were quantified using a semi-quantitative method, namely the ACD scoring system. Briefly, the dots corresponding the AAV vector genome probe were counted manually and the cells were binned into the following 5 categories: ACD score 0 corresponds to no staining or <1 dot/ 10 cells, ACD score 1 corresponds to 1 – 3 dots/cell, ACD score 2 corresponds to 4 – 9 dots/cell and none or very few dots clusters, ACD score 3 corresponds to 10 – 15 dots/cells and/or <10% dots are in clusters and ACD score 4 corresponds to >15 dots/cell and/or >10% dots are in clusters.

#### AAV transduction and treatments

iMGLs were infected with AAV6 or AAV8 at an MOI of 10^5^ for a 24 h, 48 h and 72 h, respectively. To activate the iMGLs, the cells were treated with 1 mM ATP (R0441, ThermoFisher) for 6 h or 250 ng/ml LPS (L2637, Sigma Aldrich) for 24 h. To induce γH2AX foci in iMGLs, the cells were treated with 50 µM etoposide (341205, Sigma Aldrich) for 6 h.

#### RNA extraction, DNAse treatment, cDNA preparation and qPCR analysis

To extract RNA, the iMGLs were resuspended in Qiazol and further processed with a RNA clean and concentrator kit (ZR1016, Zymo Research). The extracted RNA was quantified using a Nanodrop. Any residual contaminating DNA was eliminated by employing the Turno DNase kit according to the manufacturer’s instructions. 100 ng of DNase treated RNA was used for cDNA preparation by following the manufacturer’s instructions. The resulting cDNA was diluted 1:10 and qPCR reaction was performed using the PowerTrack SYBR Green Master Mix in a CFX96 Touch Real-Time PCR detection system. The relative gene expression values for the target genes was calculated using the Livak model of relative quantification ^57^.

#### Quantification of γH2AX foci

The mean gray value measurement of the γH2AX foci was performed by employing ImageJ (version 1.54p) as follows: In order to segment the nuclei of each cell, the DAPI was denoised and smoothened using the Gaussian Blur filter : Process -> Filters -> Gaussian Blur, with a value of sigma of 4. The object of interest (nucleus) was separated from the background by employing the Otsu thresholding technique: Image -> Adjust -> Threshold. Any background holes in the nuclei was filled using the Fill Holes command: Process -> Binary -> Fill Holes. Any isolated pixel was eliminated by using the following command: Process -> Binary -> Open. Overlapping nuclei were separated from each other using the Watershed command: Process -> Binary -> Watershed. In the Analyse Particles tab, the size option was set between 10 and infinity and any nuclei situated on the edge of the image was excluded from the analysis by checking the box ‘Exclude from edges’.

### Statistical Analysis

All the statistical analysis was performed in R (version 4.5.1) using the R package, rstatix (0.7.3). iMGL morphometric analysis was performed in a blinded manner. Statistical significance was calculated using the Kruskal-Wallis test with Dunn’s multiple comparison test. The graphs were plotted in R using the following R packages: dplyr (1.2.1), ggplot2 (4.0.2), cowplot (1.2.0), RColorBrewer (1.1-3).

